# *Phytoene synthase* modulates seed longevity via the action of β-carotene derived metabolites

**DOI:** 10.1101/2025.06.05.658166

**Authors:** Abhijit Hazra, Sombir Rao, Xuesong Zhou, Molly J Christel, Emalee Wrightstone, Yong Yang, Tara Fish, Theodore W Thannhauser, Jiping Liu, Michael Mazourek, Li Li

## Abstract

Seed longevity, the seed’s ability to stay viable over time, is an important trait in agriculture. Despite extensive research, seed longevity remains one of the fundamental topics in plant biology. Here, we discovered the novel role of carotenoid metabolites in prolonging seed lifespan. We found that *phytoene synthase* (*PSY*), the gene encoding a major rate-limiting enzyme in carotenoid biosynthesis, modulated seed storability in Arabidopsis under both natural and artificial aging conditions. Seeds from the *PSY* overexpression (OE) lines exhibited significantly enhanced lifespan with low levels of reactive oxygen species (ROS), a major factor affecting longevity, whereas those from the *psy* mutants had decreased viability with high ROS levels. While lutein and β-carotene were detected in seeds, only β-carotene and its derived apocarotenoids, *i.e.* β-cyclocitral and β-ionone, were found to improve seed lifespan. Notably, the *carotenoid cleavage dioxygenase 1* and *4* (*ccd1 ccd4*) double mutant and *PSY ccd1 ccd4* seeds showed significantly reduced seed longevity, indicating that β-carotene cleavage is necessary for or it is apocarotenoids playing the role in preserving seed lifespan. Comparative proteomic analysis identified TIP2;2, an aquaporin protein, which showed differential abundances in seeds of *PSY* OE and *psy* mutant vs wild type. The mutant *tip2;2* had reduced seed longevity, and its promoter was transactivated by apocarotenoids. Collectively, this study uncovers a novel role of apocarotenoids in protecting seed longevity and highlights the importance of seed carotenoid production in strengthening agriculture.

**One Sentence Summary:** *Phytoene synthase*, the gene encoding a major rate-limiting enzyme in carotenoid biosynthesis, modulates seed longevity via β-carotene derived apocarotenoids and an aquaporin protein TIP2;2 identified is a new player that responds to apocarotenoid signaling and influences seed longevity.

## Introduction

Seed longevity refers to seed’s ability to maintain viable over extended periods of storage. This remarkable trait is vital for plant survival, biodiversity, and agriculture. It enables seeds to withstand adverse conditions, such as drought, extreme temperatures, or long-term storage before germinating. There is considerable variation among plant species: some seeds remain viable for mere weeks whereas others can endure for centuries under optimal environmental conditions. Seed longevity largely depends on seed’s abilities to mitigate oxidative stress, maintain cellular integrity during desiccation, and repair damage upon rehydration (Sano et al. 2016; Zhou et al. 2020). Improvement of these abilities reduces deterioration and facilitates the maintenance of seed longevity. Recent studies have shed light on the molecular and physiological bases that influence seed’s ability to stay viable (Sano et al. 2016; Zhou et al. 2020; Tan et al. 2025).

The longevity of seeds at the molecular level is governed by a complex interplay of protective mechanisms that protect cellular integrity during storage (Sano et al. 2016). Key factors contributing to this process include the mechanisms to preserve DNA, proteins, and cellular membranes, all of which are susceptible to damage from oxidative stress, desiccation, and aging. Seeds accomplish these through the accumulation of protective compounds, such as antioxidants, late embryogenesis abundant proteins, and heat shock proteins, which collectively provide a safeguard against dehydration induced damage and oxidative injury (Sattler et al. 2004; Wu et al. 2011; Renard et al. 2020).

Reactive oxygen species (ROS) play a prominent role among various forms of cellular and metabolic processes involved in seed aging (Zhang et al. 2021). While functioning as signaling molecules, excess ROS accumulation during storage is detrimental to seed longevity (Bailly 2014; Jeevan Kumar et al. 2015). Elevated ROS causes oxidative damage, which leads to membrane deterioration, enzyme inactivation, and genetic damage, critical factors contributing to seed aging and loss of viability (Waterworth et al. 2010; Li et al. 2017; Oenel et al. 2017; Wiebach et al. 2020). Plants have evolved ROS scavenging systems that encompass both enzymatic and non-enzymatic antioxidants (Dumanović et al. 2021). Enzymatic defenses include catalase (CAT), ascorbate peroxidase (APX), and superoxide dismutase (SOD). Non-enzymatic antioxidants comprise molecules, such as ascorbic acid, glutathione, flavonoids, tocopherols, and carotenoids. The ROS scavenging systems work synergistically to mitigate oxidative stress and maintain ROS homeostasis within cells. Various studies have shown that alteration of the ROS detoxification systems affects seed longevity, underscoring the importance of effective ROS management for extending seed lifespan (Sattler et al. 2004; Chen et al. 2016; Wang et al. 2022).

In addition, plants have evolved various repair mechanisms to maintain seed viability (Sano et al. 2016; Zhou et al. 2020). The formation of isoaspartate at aspartate/asparagine residues during seed maturation and aging impairs protein function and reduces seed viability and vigor. Protein-L-isoaspartyl methyltransferase repairs these isoaspartate-induced damage and improves seed longevity (Ogé et al. 2008; Verma et al. 2013). Methionine sulfoxide reductases (MSR) are a class of protein-repairing enzymes that have been reported to improve seed longevity by mitigating damaged induced by methionine sulfoxide in plants (Châtelain et al. 2013; Hazra et al. 2022). DNA repair enzymes such as those in the base excision repair pathway are activated upon rehydration to rectify accumulated damage, facilitating successful germination (Waterworth et al. 2010, 2022). Despite extensive studies, the factors and mechanisms governing seed longevity remain to be fully elucidated.

Aquaporins are membrane channel proteins that play crucial roles in plant water relations and stress responses by facilitating the transport of water and small neutral molecules across cellular membranes (Maurel et al., 2008). These proteins are essential for maintaining water homeostasis, regulating hydraulic conductivity, and mediating cellular responses to various environmental stresses such as drought, salinity, and temperature extremes. Beyond water transport, some aquaporins also facilitate the movement of other small molecules like ROS, glycerol, CO, and metalloids and metal chelate, contributing to diverse physiological processes, including stress signaling, photosynthesis, and nutrient uptake (Srivastava et al. 2016; Wang et al. 2017; Handa et al. 2022).

Carotenoids like β-carotene and lutein are isoprenoids, which are vital for plants (Rodriguez-Concepcion et al. 2018; Sun et al. 2022). They are essential for photosynthesis and protect plants against photooxidative damage. Carotenoids are also potent antioxidants in plants and have been shown to effectively counter ROS and safeguard biomolecules from oxidative harm (Stahl and Sies 2003). In addition, carotenoids serve as precursors for apocarotenoids, which are generated through enzymatic cleavage by a group of carotenoid cleavage dioxygenases (CCDs) or non-enzymatic oxidation triggered by ROS. Apocarotenoids have been shown to be involved in many aspects of a plant’s life (Hou et al. 2016; Moreno et al. 2021; Beltrán and Wurtzel 2025). For example, the β-carotene derived β-cyclocitral is a stress signal to enhance oxidative stress tolerance in plants (Ramel et al. 2012; D’alessandro et al. 2018; Havaux 2020) and also possesses other known and unknown functions (Dickinson et al. 2019; Mitra et al. 2021; Deshpande and Mitra 2023; Rao et al. 2024b). Carotenoids and apocarotenoids bolster plant resilience, support reproductive success, and enable adaptation to environmental challenges through their protective and regulatory functions (Moreno et al. 2021; Beltrán and Wurtzel 2025).

In the carotenoid biosynthesis pathway, phytoene synthase (PSY) is a key rate-limiting enzyme, whose activity greatly affects carotenoid production, particularly β-carotene level in plants (Paine et al. 2005; Zhou et al. 2022). Our previous study shows that overexpression (OE) of *PSY* increases carotenoid content in Arabidopsis seeds (Sun et al. 2021). Interestingly, in this study we found that *PSY* expression modulated seed longevity under both natural and artificial aging conditions and affected seed ROS level and antioxidant enzyme activities. We demonstrated that it was the β-carotene derived apocarotenoids playing a significant role in preserving seed longevity in Arabidopsis seeds. Moreover, we identified an aquaporin protein as a new player in response to apocarotenoid signaling to contribute to seed longevity. These findings uncover a previously unrecognized function of apocarotenoids in enhancing seed storability and highlight the significance of carotenoid production in improving agricultural sustainability.

## Results

### *PSY* regulates seed longevity under both natural and artificial aging

To investigate whether *PSY* modulates seed longevity and viability, Arabidopsis wild-type (WT), Arabidopsis transgenic lines overexpressing *Solanum lycopersicum PSY2* (*PSY* #16 and #23) generated in our previous study (Cao et al. 2019), and two independent *psy* T-DNA mutant lines were used. Since Arabidopsis harbors only one copy of *PSY,* its full knockout is lethal. Thus, two *psy* lines with non-lethal phenotype were selected, which gave > 65% reduction in *PSY* transcription relative to WT (Supplementary Fig. S1).

Freshly harvested and one year naturally aged Arabidopsis seeds were germinated on ½ MS medium and counted for germination rates. All genotypes exhibited >95% germination rates with no significant differences among the freshly harvested seeds from WT, *PSY* OE and *psy* mutant lines (Fig. 1A and B). However, after one year of natural aging at room temperature, *PSY* OE seeds maintained 93-95% germination rates, WT seeds were at 86%, and *psy* mutants showed significantly reduced germination rates of 41-43% (Fig. 1A and B). Furthermore, the young seedlings from freshly harvested seeds germinated at the same time and grew similarly (Supplementary Fig. S2). After 1 year of aging, the *PSY* OE lines grew faster with longer roots, whereas *psy* mutants had shorter roots than WT, showing differential levels of seedling vigor (Supplementary Fig. S3A).

**Figure 1.**
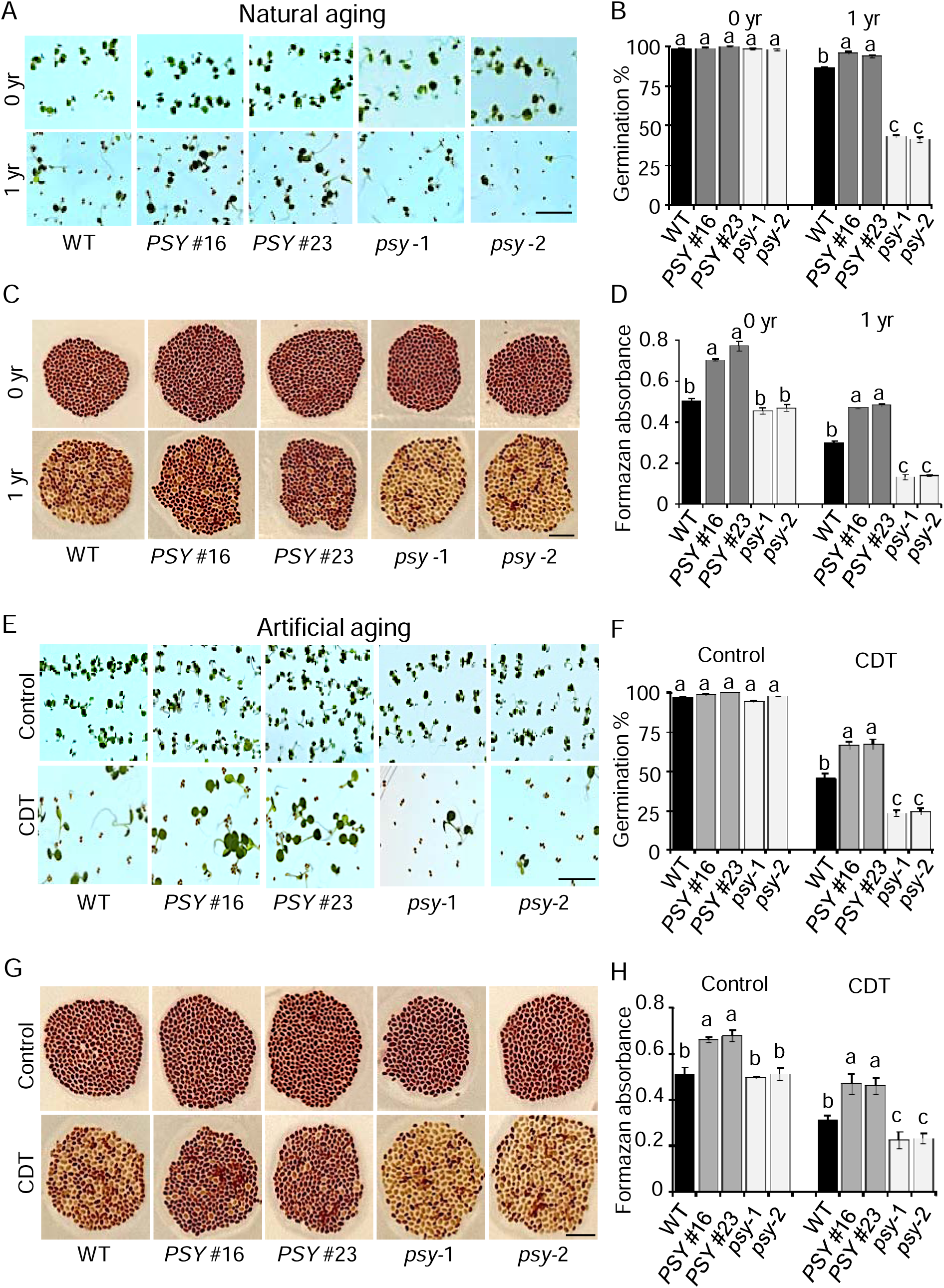
*PSY* expression affects seed longevity during natural and artificial aging. All genotype seeds were harvested on the same day. For natural aging, seeds were stored at room temperature in sealed bags for 1 year. **A)** Images of germinated seeds of different genotypes after 72 hrs post radicle emergence before and after natural aging. Upper and low panels show freshly harvested (0 yr) and after one year (1 yr) storage, respectively. WT, wild type. *PSY* overexpression (OE) lines (#16 and #23) (Cao et al., 2019), and two *psy* T-DNA insertion mutants. **B)** Germination percentages of different genotypes before and after one-year natural aging. **C)** Tetrazolium staining for seed viability. Dark color indicates more viable seed tissues. **D)** Formazan absorbance of WT, *PSY* OE and *psy* seeds before and after natural aging. **E)** Images of germinated seeds of different genotypes after 72 hrs post radicle emergence in control and after artificial aging. For artificial aging, a controlled deterioration test (CDT) was conducted by treating the seeds at 45 °C with 100% relative humidity for 4 days prior to germination on ½ MS medium. **F)** Germination percentages of different genotypes in control and after CDT treatment. **G)** Tetrazolium staining for seed viability in control and after CDT treatment. **H)** Formazan absorbance of WT, *PSY* OE and *psy* seeds. Scale bars 0.5 cm. Data in **B**, **D**, **F**, and **H** are means ± SD of three biological repeats. Different letters indicate significant differences in means determined using one-way ANOVA with Duncan’s multiple range test (*p* = 0.01).

A tetrazolium assay, frequently used to test seed viability (Berridge et al. 2005), was performed to confirm further the longevity difference observed among these genotypes. Freshly harvested seeds of WT, *PSY* OE lines, and *psy* mutants were almost uniformly stained in dark red due to the formation of formazan from the reduction of tetrazolium salt by living seeds. In contrast, after 1 year of natural aging, *PSY* OE seeds exhibited darker stain than WT, and the *psy* mutant seeds showed reduced color intensity with less viable seeds (Fig. 1C). When the formazan production was quantified, *PSY* OE seeds exhibited significantly higher and the *psy* mutant seeds substantially lower levels than WT seeds after natural aging (Fig. 1D), showing variation of viable seeds in the different genotypes. The *PSY* OE seeds were also found to have significantly higher formazan production than WT in the freshly harvested seeds (Fig. 1D).

To further assess the role of *PSY* in seed longevity, we performed a control deterioration test (CDT), an accelerated aging or artificial aging assay that has been shown to mimic natural aging with similar molecular events (Rajjou and Debeaujon 2008). Fresh harvested seeds were exposed to 45°C and 100% relative humidity (RH) for 4 days prior to germination tests. As shown above without CDT treatment, no significant difference in the germination rates was observed among seeds from WT, *PSY* OE lines, and *psy* mutants, where all genotypes had >95% germination (Fig. 1E and F). After artificial aging treatment, *PSY* OE seeds showed higher germination rates than WT, whereas *psy* mutants had significantly reduced rates (Fig. 1E and F). The *PSY* OE lines also grew faster, and the *psy* mutants grew slower than WT, showing differential levels of seedling vigor (Supplementary Fig. S3B). Seed viability analyzed by tetrazolium staining and formazan production showed strong correlations with these genotypes. The *PSY* OE lines exhibited significantly higher viability, whereas *psy* mutants showed markedly lower viability compared to WT after CDT treatment (Fig. 1G and H).

To validate these findings in another species, we also analyzed tomato wild-type (AC) and *psy1* mutant seeds (Fray and Grierson 1993). The freshly harvested AC and *psy1* mutant seeds had similar germination rates of 97-98% (Supplementary Fig. S4A and B). However, after one year of storage, the *psy1* mutant seeds showed a greatly reduced germination rate, much lower than the AC control (Supplementary Fig. S4A and B). We compared seed viability using the tetrazolium staining and found that both freshly harvested AC and *psy1* mutant seeds were uniformly stained. However, after one year of storage, the *psy1* mutant seeds showed significantly weaker staining than AC (Supplementary Fig. S4C). Quantification of the formazan levels revealed that *psy1* seeds had a significantly lower formazan production than AC (Supplementary Fig. S4C). Similar trends were also observed after artificial aging (Supplementary Fig. S4E-H).

Together, these findings showed that *PSY* expression directly affects seed longevity under both natural and artificial aging. Elevated *PSY* expression improves seed lifespan, while *PSY* deficiency leads to accelerated deterioration during aging.

### *PSY* expression affects ROS levels in Arabidopsis seeds

ROS can act as a signaling molecule to promote dormancy breaking, playing a significant role in seed physiology (Li et al. 2022). However, elevated ROS levels compromise seed viability and overall seed health (Bailly 2014; Jeevan Kumar et al. 2015). To investigate whether the seed longevity differences observed among the various Arabidopsis genotypes are correlated to ROS levels, we examined hydrogen peroxide (H_2_O_2_) accumulation in the seeds of WT, *PSY* OE and *psy* mutants with and without artificial aging treatments. DAB (3,3’-diaminobenzidine) staining detects ROS, particularly H_2_O_2_. *PSY* OE seeds stained lighter and *psy* mutant seeds stained darker than WT particularly after CDT treatment (Fig. 2A), showing a direct correlation with *PSY* expression levels. We then quantified H_2_O_2_ levels in seeds of these genotypes. Seeds of *PSY* OE lines accumulated significantly lower levels of H_2_O_2_ and the *psy* mutants had substantially higher contents than WT with and without CDT treatments (Fig. 2B). In addition, we quantified the levels of malondialdehyde (MDA) that reflects lipid peroxidation (Draper and Hadley 1990) and influences seed longevity and vigor. MDA levels were also significantly lower in the *PSY* OE seeds and higher in the *psy* mutants compared with WT (Fig. 2C).

**Figure 2.**
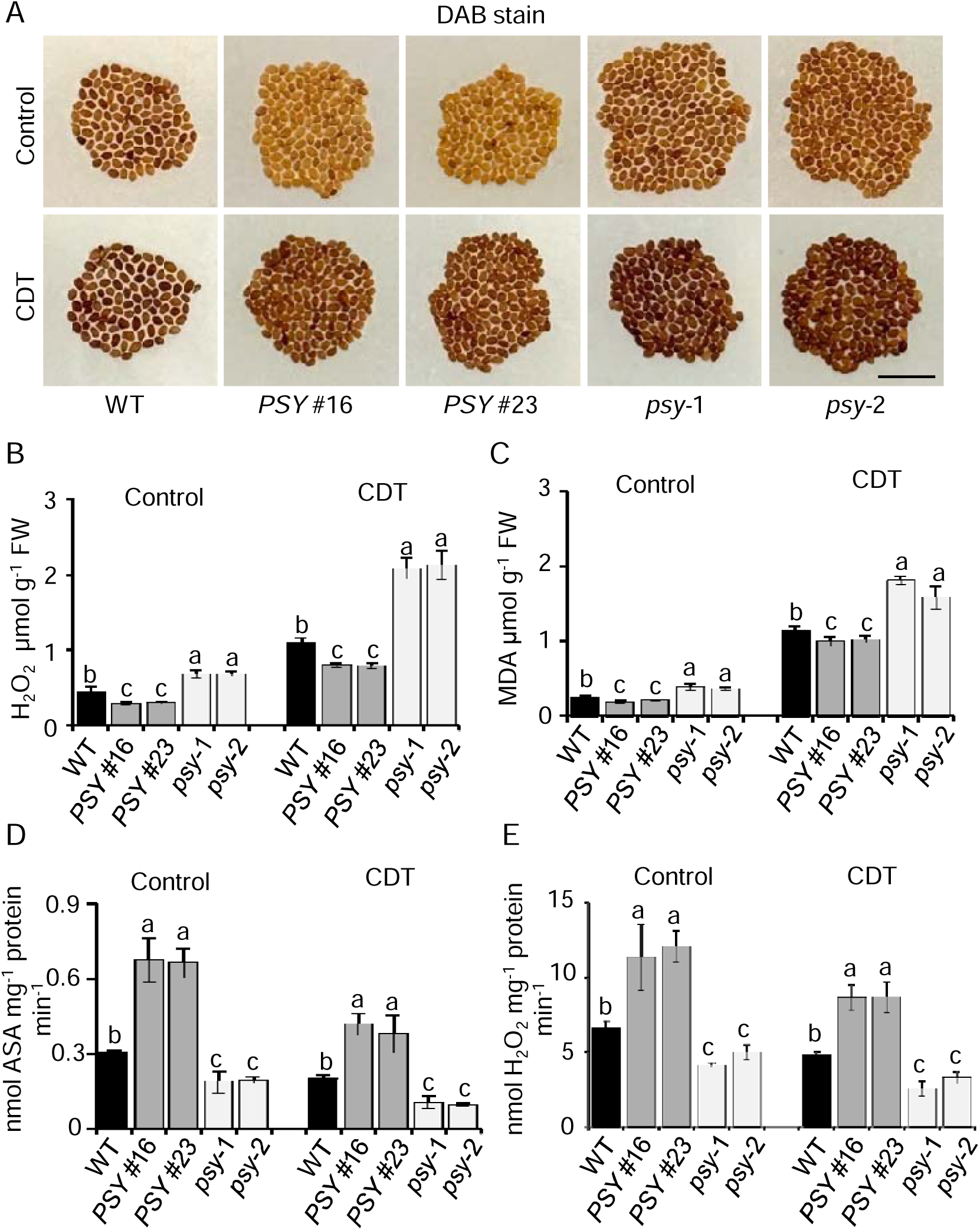
*PSY* expression modulates ROS levels in seeds. **A)** 3,3’-diaminobenzidine (DAB) staining shows differential accumulation of H_2_O_2_ in wild-type (WT), *PSY* OE and *psy* mutant seeds in control and after controlled deterioration test (CDT) treatment. Dark staining of the seeds indicates more H_2_O_2_ accumulation. Scale bars 0.5 cm. **B)** Quantitative analysis of H_2_O_2_ in seeds of various genotypes in control and after artificial aging. **C)** Lipid peroxidation analysis by quantifying malondialdehyde (MDA) in seeds of various genotypes. **D**) Ascorbate peroxidase (ASA) and **E)** catalase activity in seeds of various genotypes in control and after CDT treatment. Data in B-E are means ± SD of three biological repeats. Different letters indicate significant differences in means determined using one-way ANOVA with Duncan’s multiple range test (*p* = 0.01).

It is well known that antioxidative machinery plays a pivotal role in regulating ROS homeostasis (Bailly., 2014). We compared the enzymatic activity of ascorbate peroxidase (APX) and catalase (CAT) in seeds of WT, *PSY* OE, and *psy* mutant lines. APX activity was significantly higher in the *PSY* OE seeds and substantially lower in the *psy* mutants than the WT seeds both with and without CDT treatments (Fig. 2D). Similarly, catalase activity also showed the same trends across these genotypes (Fig. 2E).

In addition, we analyzed the transcript levels of *AtAPX1*, *AtAPX3*, *AtAPX6, AtCAT2*, and *AtCAT3* in seeds of WT, *PSY* OE, and *psy* mutant lines. These genes expressed at significantly higher levels in the *PSY* OE seeds and lower levels in the *psy* mutant seeds than WT (Supplementary Fig. S5). Together, these results showed that *PSY* expression influences ROS production and antioxidative enzymes in affecting seed longevity.

### Carotenoid metabolites modulate seed longevity

*PSY* OE has been shown to increase carotenoid levels in Arabidopsis seeds (Sun et al. 2021). To examine whether the observed seed longevity differences among various genotypes correlate with carotenoid metabolites produced, the carotenoid content and composition in seeds of these lines with and without artificial aging treatments were analyzed. *PSY* OE seeds contained significantly higher levels of total carotenoids, whereas *psy* mutant seeds had substantially lower levels than WT seeds both in the control and after artificial aging treatment (Fig. 3A). Noticeably, the carotenoid levels of all genotypes exhibited dramatic reductions after the artificial aging treatment for 4 days, consistent with the observation that carotenoids are very heat labile in Arabidopsis seeds (Sun et al. 2021). After the aging treatment, the PSY protein level was also dramatically reduced (Supplementary Fig. S6).

**Figure 3.**
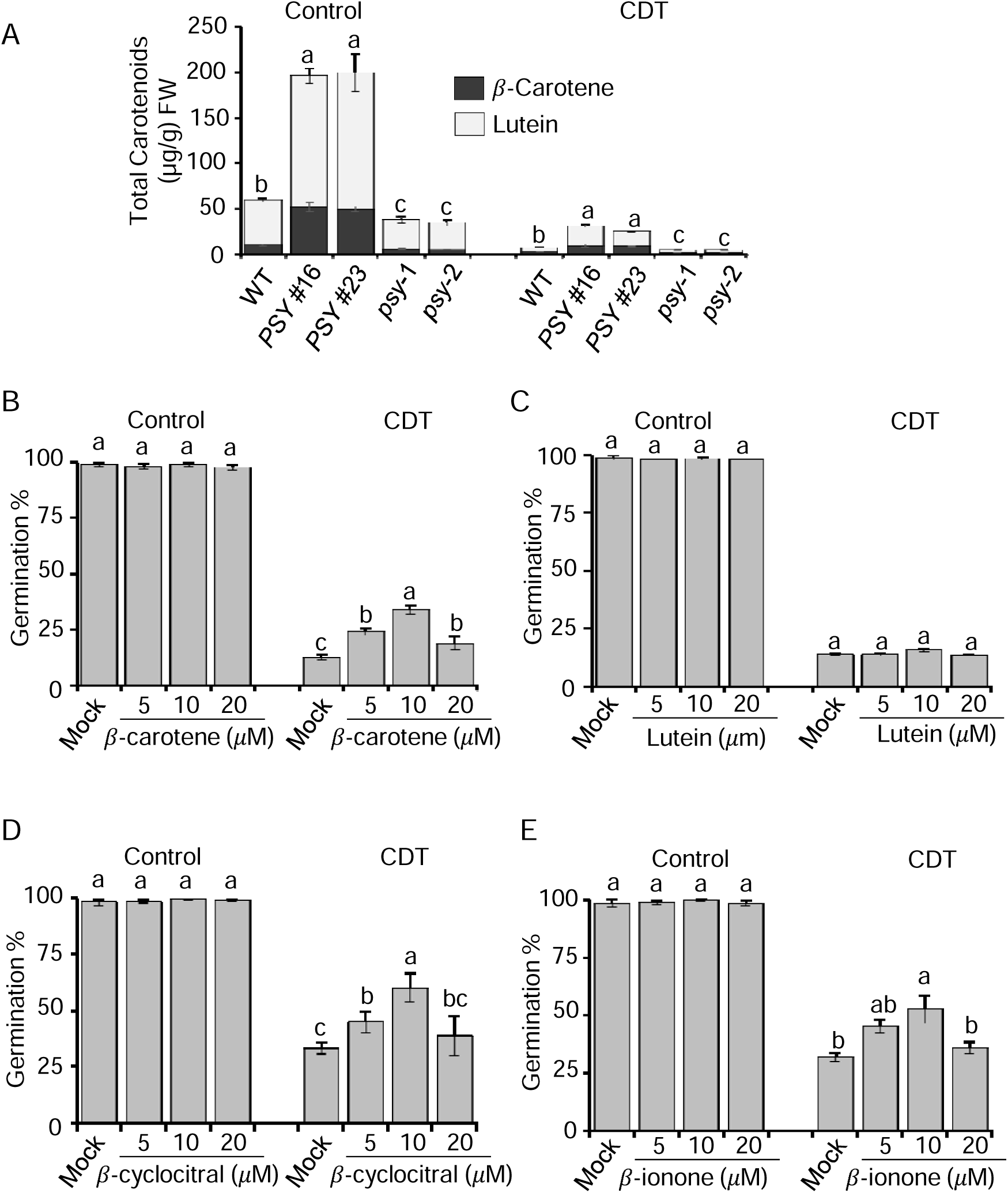
Carotenoid metabolites regulate seed longevity in Arabidopsis. **A)** Carotenoid levels in seeds of wild-type (WT), *PSY* OE, and *psy* mutant lines without (control) and after artificial aging (CDT) treatment, analyzed by UCP2. FW, fresh weight. **B-E)** Germination percentages of WT seeds pretreated with **B)** *β*-carotene, **C)** lutein, **D)** *β*-cyclocitral, and **E)** *β*-ionone at 5 *μ*M, 10 *μ*M, 20 *μ*M along with mock for 3 hours, followed by without (control) and with CDT treatment for 4 days prior to germination on½ MS medium. All data are means ± SD of three biological repeats. Different letters indicate significant differences in means determined using one-way ANOVA with Duncan’s multiple range test (*p* = 0.01).

Lutein and β-carotene were detected in Arabidopsis seeds, whose levels were greatly increased in the *PSY* OE seeds and decreased in *psy* mutant seeds (Fig. 3A). To check whether β-carotene or lutein imparts seed longevity, we treated WT Arabidopsis seeds with various concentrations (0, 5, 10, and 20 μM) of β-carotene and lutein prior to aging treatment. Without CDT treatment, seeds exposed to either β-carotene or lutein had similar germination rates of >95% as mock control (Fig. 3B and C). After CDT treatment, enhanced seed germination rates were observed with increased doses of β-carotene in comparison with mock control, where a β-carotene supplement at 10 μM showed the highest increase of germination rate (Fig. 3B). In contrast, the lutein treatment showed no improvement on seed germination compared to mock (Fig. 3C). These results indicate a specific role of β-carotene in improving seed longevity.

β-carotene is a precursor of various bioactive apocarotenoids (Moreno et al. 2021). It was intriguing to check whether the β-carotene derived apocarotenoids can also enhance seed longevity. β-cyclocitral and β-ionone are the two most studied apocarotenoids derived from β-carotene (Havaux 2020; Moreno et al. 2021). Thus, we performed similar experiments in WT Arabidopsis seeds, which treated with and without β-cyclocitral and β-ionone at various concentrations. Both β-cyclocitral and β-ionone-treated seeds improved seed longevity with strongest effect observed at 10 μM after CDT treatment (Fig. 3D and E). These findings suggest that the increased seed longevity observed in *PSY* OE lines likely results from increased production of β-carotene and/or its derived apocarotenoids.

### Apocarotenoids are the key regulatory elements that modulate seed longevity

To investigate whether β-carotene or the apocarotenoid molecules are important in imparting seed longevity, we first examined the *ccd1 ccd4* (*ccd1/4*) double mutant and generated Arabidopsis transgenic lines that overexpressed *PSY* in the double mutant background (*PSY ccd1/4*). Three homozygous transgenic lines with high PSY protein levels in *ccd1/4* mutant background were selected (Fig. 4A; Supplementary Fig. S7). Their germination rates were examined. In the control without CDT treatment, all genotypes showed similarly high germination rates of >95%. After artificial aging, while the *PSY* OE seeds showed significantly higher germination rate than WT as shown above, the *ccd1/4* mutant seeds had more than half reduced germination rate compared to WT (Fig. 4B, Supplementary Fig. S8A), indicating that CCD activity greatly affects seed longevity. Furthermore, when *PSY* was overexpressed in the *ccd1/4* mutant, the *PSY ccd1/4* lines had similar low germination rates as the *ccd1/4* seeds (Fig. 4B, Supplementary Fig. S8A), indicating that carotenoid cleavage into apocarotenoids is required for the *PSY* regulated seed longevity. These data suggest the definitive role of apocarotenoids in modulating seed longevity in Arabidopsis seeds.

**Figure 4.**
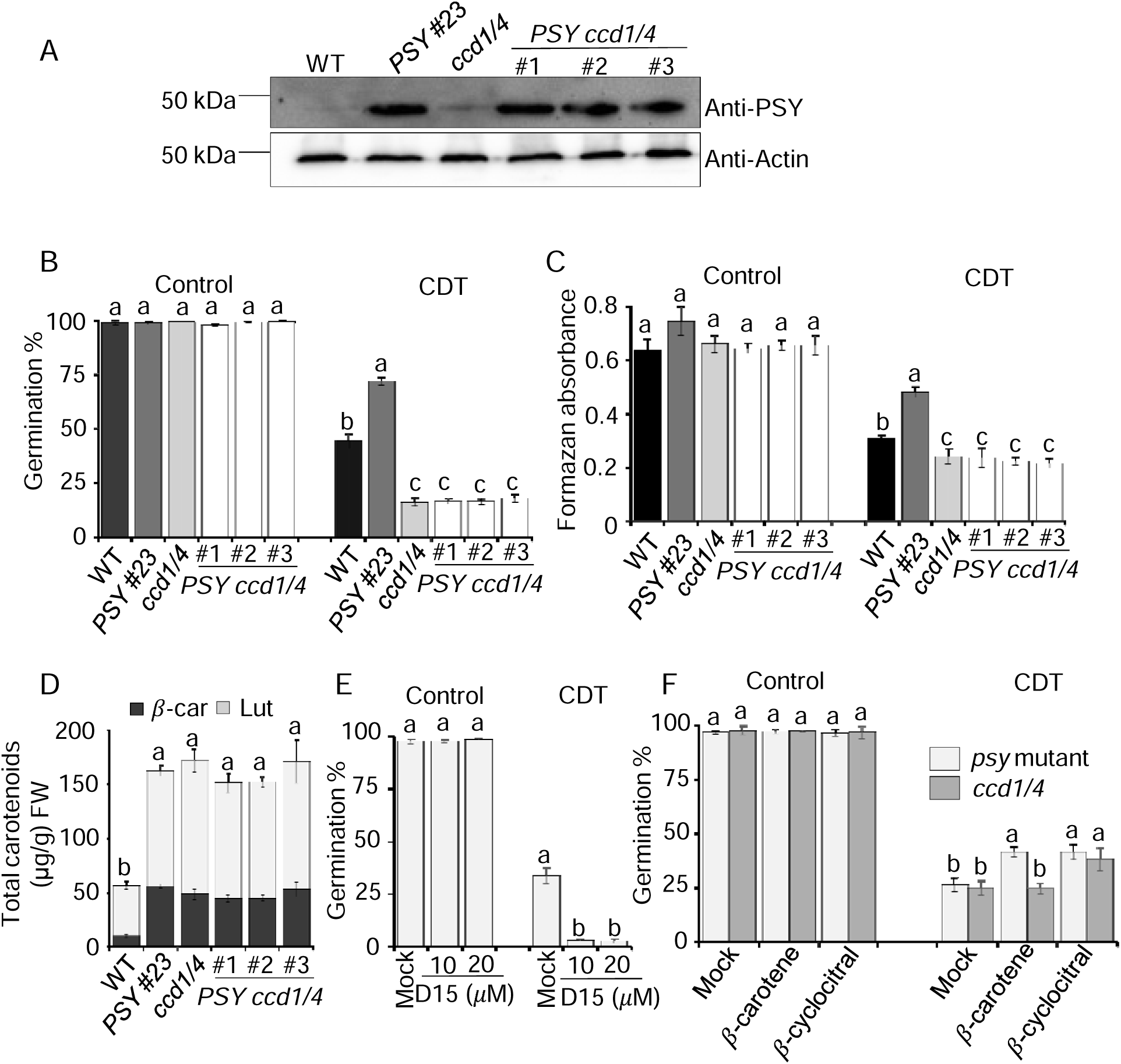
Apocarotenoids play a key role in maintaining seed longevity. **A)** Immunoblot analysis of PSY protein levels using anti-PSY antibody from 25 *μ*g seed proteins of WT, *PSY* OE (#23), *ccd1/4* double mutant, and three *PSY ccd1/4* lines. Anti-actin blot was used for a loading control (lower panel). **B)** Germination percentages of freshly harvested seeds of various genotypes in control and after controlled deterioration test (CDT) treatment. **C)** Quantification of formazan absorbance after tetrazolium staining. **D)** Carotenoid levels in seeds of various genotypes analyzed by UCP^2^. FW, fresh weight; *β*-Car, *β*-carotene; Lut, lutein. **E)** Germination rates of wild-type seeds treated with D15 at 10 *μ*M and 20 *μ*M for 3 hours along with mock followed by with and without CDT aging treatment. **F)** Germination rates of *psy* and *ccd1/4* mutant seeds treated with *β*-carotene and *β*-cyclocitral along with mock followed by with and without CDT aging treatment. All bar data are means ± SD of three biological repeats. Different letters indicate significant differences in means determined using one-way ANOVA with Duncan’s multiple range test (*p* = 0.01).

Seed viability was also examined by tetrazolium staining and quantified by measuring formazan production. Without artificial aging, the tetrazolium staining and formazan absorbance showed no much differences among all genotypes (Fig. 4C, Supplementary Fig. S8B). However, after artificial aging, the *ccd1/4* mutant seeds exhibited less tetrazolium staining and significantly lower formazan absorbance, whereas the *PSY* OE seeds showed significantly higher compared to WT (Fig. 4C, Supplementary Fig. S8B). Moreover, seeds of the *PSY ccd1/4* lines also did not increase the formazan production compared to *ccd1/4* mutant seeds (Fig. 4C), further supporting the role of apocarotenoids in affecting seed viability after artificial aging.

To validate that β-carotene itself contributes minimal in seed longevity, we analyzed the carotenoid levels. The seeds of *PSY* OE, *ccd1/4*, and *PSY ccd1/4* lines all accumulate significantly higher β-carotene levels than WT (Fig. 4D). Even *ccd1/4* and *PSY ccd1/4* seeds showed greatly reduced seed longevity, they contained similar levels of β-carotene as *PSY* OE seeds. These results support that it is not β-carotene but its derived apocarotenoids in modulating seed longevity.

To confirm the role of apocarotenoids, we also treated WT seeds with D15, an inhibitor of CCDs that suppresses apocarotenoid production (Van Norman et al. 2014). Without artificial aging, there were no significant changes in germination rates between the seeds treated with or without D15 (Fig. 4E, Supplemental Fig. S9A). However, following CDT treatment, the germination rates for seeds treated with D15 significantly decreased to under 10%, much lower than mock with an over 30% germination rate (Fig. 4E, Supplemental Fig. S9A). These results indicate the importance of CCD catalyzed apocarotenoid production in regulating seed longevity.

To further support the role of apocarotenoids, we also treated *psy* and *ccd1/4* mutant seeds with 10 µM β-carotene and 10 µM β-cyclocitral. Before artificial aging, there were no significant differences in germination rates among the genotypes following metabolite treatments. However, after CDT, β-carotene treatment only improved the germination percentage of *psy* mutant, but not of *ccd1/4* mutant in comparison with mock controls (Fig. 4F), showing the importance of β-carotene conversion into apocarotenoids by CCDs. In contrast, β-cyclocitral treatment was found to enhance seed longevity in both *psy* and *ccd1/4* mutants compared to the mocks (Fig. 4F). Consistently, when natural aging *psy-1* and *psy-2* seeds were treated with 10 µM β-cyclocitral, significantly increased germination rates were observed (Supplemental Fig. S9B). These results further emphasize the specific role of apocarotenoids but not β-carotene directly in modulating seed longevity.

### ROS homeostasis is regulated through apocarotenoids but not carotenoids during seed aging

Carotenoids are endowed with antioxidant properties. To examine whether carotenoids themselves as antioxidants contribute to the *PSY* regulated ROS levels during seed aging, we examined H_2_O_2_ accumulation, MDA level, and antioxidant enzyme activities in the seeds of *ccd1/4* and *PSY ccd1/4* lines that accumulated more carotenoids (Fig. 4D) along with WT and a *PSY* OE line. The *PSY* OE seeds showed consistent trends compared with WT as observed in Figure 2, with less ROS production and higher antioxidant enzyme activities both in control and after CDT treatment (Fig. 5). Interestingly, seeds from both *ccd1/4* and *PSY ccd1/4* lines had darker DAB staining and higher H_2_O_2_ levels than WT both in control and after CDT treatments (Fig. 5A and B). Similarly, their MDA levels were also significantly higher than WT (Fig. 5C). These results indicate that the increased carotenoid levels in *PSY ccd1/4* seeds have less impact in reducing ROS levels during aging.

**Figure 5.**
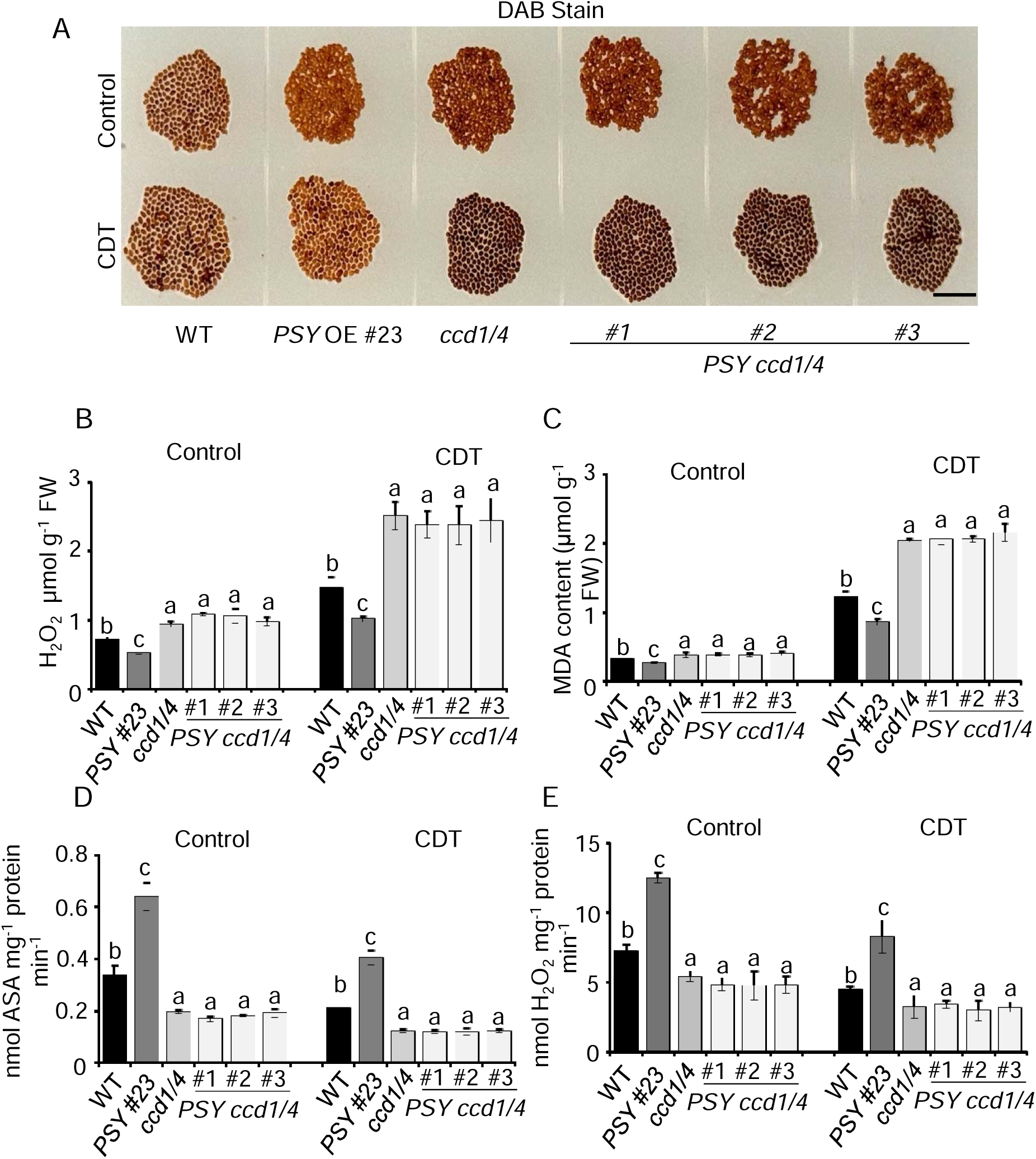
*PSY* expression modulate ROS levels through apocarotenoids but not *β*-carotene in Arabidopsis seeds upon artificial aging. **A)** 3,3’-diaminobenzidine (DAB) staining shows the differential accumulation of hydrogen peroxide (H_2_O_2_) in wild-type (WT), *PSY* OE, *ccd1/4* and *PSY ccd1/4* seeds in control and after controlled deterioration test (CDT) treatment. Dark staining of the seeds indicates more H_2_O_2_ accumulation. Scale bars 0.5 cm. **B)** Quantitative analysis of H_2_O_2_ in seeds of various genotypes in control and after artificial aging. **C)** Lipid peroxidation analysis by quantifying malondialdehyde (MDA) in seeds of various genotypes. **D)** Ascorbate peroxidase (ASA) and **E)** catalase activity in seeds of various genotypes in control and after CDT treatment. Data in **B-E** are means ± SD of three biological repeats. Different letters indicate significant differences in means determined using one-way ANOVA with Duncan’s multiple range test (*p* = 0.01).

We also examined the APX and CAT enzymatic activities in the seeds of *ccd1/4* and *PSY ccd1/4* lines. APX activity was significantly lower in the seeds of *ccd1/4* and *PSY ccd1/4* lines than the WT seeds both with and without CDT treatments (Fig. 5D). Similarly, catalase activity also showed the same trends across these genotypes (Fig. 5E). Collectively, the genetic studies reveal that the *PSY* altered ROS levels and antioxidant enzyme activities in the seeds of these genotypes are unlikely to be associated with carotenoid antioxidant properties to affect *PSY*-enhanced seed longevity.

### TIP2;2, an aquaporin protein, plays a novel role in regulating seed longevity

Plants have developed multiple strategies to maintain seed longevity for their survival and reproductive success (Sano et al. 2016). To identify new factors that potentially play a role in seed longevity, a comparative proteomic analysis was performed on seeds of Arabidopsis. A total of 4,802 seed proteins were identified from three biological replicates of Arabidopsis WT, *PSY* OE, and *psy* mutant seeds (Supplementary Table S1). Among them, 180 proteins were upregulated when compared *PSY* OE with WT seed proteins (≥ 1.5 fold change) and 171 proteins were downregulated when compared *psy* mutant with WT seed proteins (≤ 0.5 fold change) (Fig. 6A; Supplementary Table S2). Tonoplast intrinsic proteins (TIPs) are known to contribute to ROS detoxication within cells (Bienert et al. 2007; Wang et al. 2014). Notably, AT4G17340 (TIP2;2) was identified as an overlapping protein with a highest abundance in *PSY* OE seeds and lower abundance in the *psy* mutant seeds compared to WT (Fig. 6A). Consistent with these findings, RT-qPCR analysis revealed that *TIP2;2* was expressed significantly higher in seeds of *PSY* OE and lower in *psy* mutants than WT (Fig. 6B).

**Figure 6.**
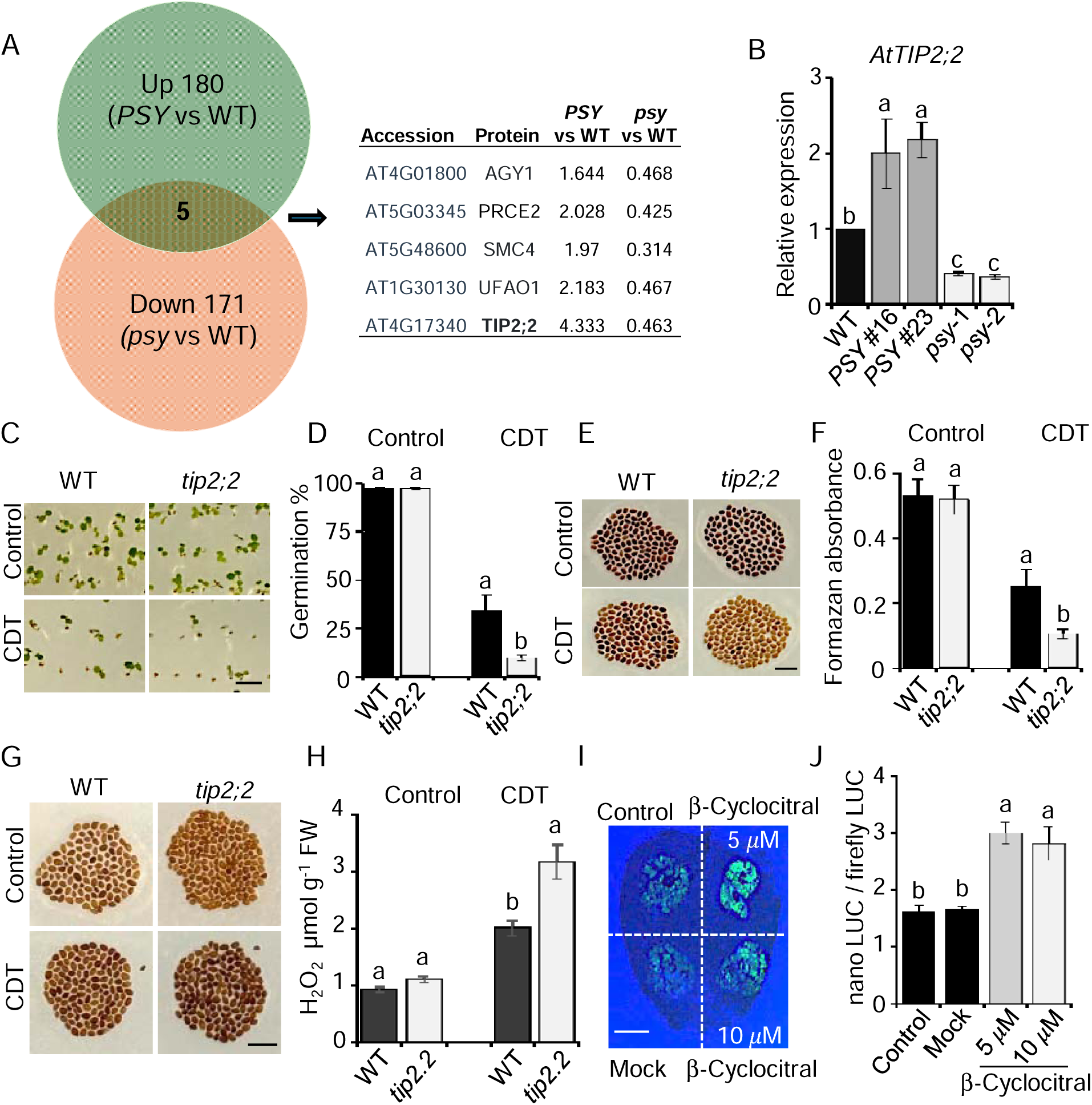
TIP2.2 protein contributes to seed longevity. **A)** Venn diagram of differential abundant proteins upregulated in *PSY* OE and downregulated in *psy* mutant seeds comparing with wild-type detected from comparative proteomic analysis. TIP2 was among the five common proteins that were up highest in *PSY* OE and down in *psy* mutant. **B)** Relative abundance of *TIP2* transcript in seeds of WT, *PSY* OE, and *psy* mutant lines. **C)** Images of germinated seeds of WT and *tip2* mutant in control and after CDT treatment. **D)** Germination rates of WT and *tip2* mutant seeds in control and after CDT treatment. **E)** Tetrazolium staining. **F)** Formazan absorbance after tetrazolium staining of WT and *tip2* mutant seeds. **G)** 3,3’-diaminobenzidine (DAB) staining in seeds of WT and *tip2* mutant in control and after CDT treatment. Dark staining of the seeds indicates H_2_O_2_ accumulation. **H)** Quantitative analysis of H_2_O_2_ levels in seeds. **I)** Trans-activation assay luciferase signals. Dual-luciferase construct driven by 2kb *Tip2.2* promoter with nanoLUC was used as a reporter and FireflyLUC under UBQ10 promoter as control (Sun et al., 2025). Transactivation of *Tip2.2* promoter with *β*-cyclocitral at 5 μM and 10 μM. **J)** Quantification of the transactivation activity. Scale bars 0.5 cm. All bar figure data are means ± SD of three biological repeats. Different letters indicate significant differences in means determined using one-way ANOVA with Duncan’s multiple range test (*p* = 0.01).

To investigate whether TIP2;2 is involved in modulating seed longevity, we compared the seed germination rate of *tip2;2* T-DNA insertion mutant, which had dramatically reduced *TIP2;2* expression (Supplementary Fig. S10), in the control and after artificial aging treatment. While both WT and *tip2;2* seeds had similar germination rates (>95%) without CDT treatment, the *tip2;2* mutant seeds showed a significantly lower germination than WT after artificial aging (Fig. 6C and D). When they were stained with tetrazolium and quantified for formazan levels, the *tip2;2* mutant seeds stained lighter with less viable seed tissues and had lower formazan production compared to WT after artificial aging (Fig. 6E and F), showing low seed viability.

TIP proteins are known to facilitate H_2_O_2_ transport from the cytosol into the vacuole for ROS detoxification in the cell (Bienert et al. 2007). We also performed ROS staining and found that the *tip2;2* mutant seeds stained darker than WT, particularly after artificial aging (Fig. 5G). Moreover, the *tip2;2* seeds accumulated more H_2_O_2_ compared to WT following CDT treatment (Fig. 5H), indicating that TIP2;2 affects ROS production during aging.

Given the differential abundance of TIP2;2 in WT, *PSY* OE, and *psy* mutant seeds and its association with seed longevity, we tested whether apocarotenoids modulate *TIP2;2* expression. We conducted a transactivation assay *in planta* using a dual-luciferase system (Sun et al. 2025). The *TIP2;2* promoter-driven luciferase existed stronger signals than controls when treated with 5 μM and 10 μM of β-cyclocitral (Fig. 5I). Quantification of the promoter activity indicated that β-cyclocitral significantly enhanced the transactivation activity of the *TIP2;2* promoter compared to non-treated and mock controls (Fig. 5J). We also treated with β-ionone that was shown to regulate seed longevity after artificial aging (Fig. 3E). Consistently, β-ionone at 5 μM and 10 μM also transactivated the *TIP2;2* promoter-driven luciferase with increased signals and activities in comparison with controls (Supplementary Fig. S10). These results indicate that *TIP2:2* promoter responds to β-cyclocitral and β-ionone signaling, although the precise mechanism by which these apocarotenoids modulate the *TIP2;2* promoter remains to be determined.

## Discussions

Seed longevity is a key agronomic trait in the domains of agriculture, seed preservation, and ecological sustainability (Sano et al. 2016; Zhou et al. 2020). Carotenoids are found in all seeds, indicating their important roles in seed biology. In this study, we discovered the novel functions of carotenoid metabolites in prolonging seed lifespan. We showed that *PSY* expression modulated seed longevity under both natural and artificial aging conditions, altered ROS levels, and influenced ROS salvaging enzyme activities. We demonstrated that it was the β-carotene derived apocarotenoids, such as β-cyclocitral and β-ionone, in preserving the seed longevity of Arabidopsis seeds. We also identified an aquaporin protein as a novel player in response to apocarotenoid signaling to modulate seed longevity. These findings enhance our understanding of metabolite-regulated seed longevity and support the critical role of carotenoid production for agriculture.

### *PSY* is an important genetic element for seed longevity

PSY is a major rate-limiting enzyme in the carotenoid biosynthetic pathway in plants (Paine et al. 2005; Zhou et al. 2022). Manipulation of *PSY* transcription has been shown to greatly affect carotenoid levels, particularly β-carotene in various tissues of many plants (Fraser et al. 2000; Paine et al. 2005; Maass et al. 2009; Naqvi et al. 2009; Zhou et al. 2015; Cao et al. 2019; Sun et al. 2021). Consistent with these studies, alteration of *PSY* expression changed carotenoid levels in seeds of Arabidopsis (Fig. 3A), showing increased carotenoid levels in the *PSY* OE lines and reduced content in the *psy* mutants. Excitingly, we discovered that *PSY* also played a novel role in seed longevity under both natural and artificial aging conditions (Fig. 1), demonstrating *PSY* as an important genetic element in regulating seed longevity.

Recent studies report that overexpression of carotenoid pathway genes not only increases carotenoid content but also enhances crop performance beyond their classical functions. Overexpression of *lycopene* β*-cyclase (LCYB*), encoding an enzyme that cyclizes lycopene, was found to promote fast growth, larger biomass, and early flowering as well as enhanced tolerance to salt and ROS stresses in tobacco (Moreno et al. 2016, 2021; Kössler et al. 2021). *LCYB* OE was also noted to increase seed yield, stress tolerance and shelf life in tomato (Mi et al. 2022). Our recent work reveals that tomato *PSY1* affects pollen fertility by regulating ROS homeostasis (Rao et al. 2024b). Here we found that the *PSY* OE seeds exhibited lower ROS accumulation and the *psy* mutants had higher levels than WT, whereas the activities of ROS-scavenging enzymes were higher in the *PSY* OE lines and lower in the *psy* mutants after artificial aging (Fig. 2). These findings strongly correlated *PSY* expression with changes of plant redox systems for seed longevity.

Oxidative stress is one of the primary contributors to seed deterioration. During storage, ROS accumulation causes lipid peroxidation, protein carbonylation, and DNA damage, which erode seed viability (Waterworth et al. 2010; Li et al. 2017; Oenel et al. 2017; Wiebach et al. 2020). Carotenoids as antioxidants have been shown to safeguard against lipid peroxidation in membranes and enhance membrane stability by quenching ROS (Havaux 1998; Young and Lowe 2001). As such, plants with higher carotenoid content often exhibit greater tolerance to oxidative damage resulting from environmental stresses (Davison et al. 2002). However, our genetic studies with *ccd1/4* and *PSY ccd1/4* lines revealed that although their seeds contained higher levels of carotenoids than WT, they exhibited much higher ROS and MDA levels than WT and *PSY* OE lines (Fig. 5), indicating that the increased carotenoids in *ccd1/4* background exhibited minimal effects on reducing ROS and MDA content. The results also suggest that the *PSY* regulated seed longevity with altered ROS levels is not associated with the carotenoids’ antioxidant properties.

The antioxidant gene expression and enzyme activity were directly correlated with *PSY* expression in the *PSY* OE and *psy* mutant seeds (Fig. 2). ROS can activate redox-sensitive genes to regulate the expression of antioxidant enzymes (Locato et al. 2018; Hong et al. 2024). Thus, *PSY* probably affects seed longevity via carotenoids derived metabolites, which act as signaling molecules to regulate redox systems.

### β-carotene derived apocarotenoids exert a novel function in seed longevity

Newly identified apocarotenoids are emerging as important signaling and regulatory molecules with new functions in plant growth and development, stress response, and communications (Havaux 2020; Moreno et al. 2021; Sierra et al. 2022; Beltrán and Wurtzel 2025). We uncovered a new role of apocarotenoids in seed longevity.

The carotenoid profile analysis in seeds of *PSY* OE, *psy* mutants, and WT detected β-carotene and lutein accumulated in Arabidopsis seeds. Evidence from chemical, molecular, and genetic analyses revealed that it was not β-carotene or lutein but the β-carotene derived apocarotenoids that play a key role in regulating seed longevity (Fig. 3 and 4). When seeds were treated with β-carotene and lutein, only the β-carotene treated seeds exhibited increased seed longevity following artificial aging, whereas lutein had no effect. β-carotene can be enzymatically cleaved by some CCD enzymes and *ccd4* mutant accumulates β-carotene in seeds (Gonzalez-Jorge et al. 2013). Although *ccd1/4* seeds contained higher level of β-carotene than WT, they did not exhibit enhanced seed longevity after artificial aging.

Furthermore, when *PSY* was overexpressed in *ccd1/4* mutant, the increased β-carotene levels had no effect on seed longevity, supporting the conclusion that β-carotene explicitly is not responsible. Instead, β-carotene derived apocarotenoids, β-cyclocitral and β-ionone, were found to effectively enhance seed longevity. Additionally, D15, an inhibitors of CCD activities (Van Norman et al. 2014b; Dickinson et al. 2019), dramatically lowered seed longevity after artificial aging. These results provide strong evidence of the involvement of β-carotene derived apocarotenoids in improving seed longevity.

β-Cyclocitral has been shown to exert various functions in plants such as in promoting tolerance to photooxidative stress (Ramel et al. 2012; D’alessandro et al. 2018) regulating root growth (Dickinson et al. 2019), modulating pollen fitness (Rao et al. 2024b), increasing defense against herbivore damage (Mitra et al. 2021), and enhancing drought and salt tolerance (D’alessandro et al. 2018). Interestingly, β-cyclocitral has been shown to promote detoxification and antioxidant enzyme activities to enhance stress tolerance (Ramel et al. 2012; Deshpande and Mitra 2023). Although less is known about β-ionone’s functions in plants compared to β-cyclocitral, it has been reported to influence plant–insect interactions and to enhance Arabidopsis resistance to fungal pathogens (Felemban et al. 2024). Together, β-cyclocitral and β-ionone along with other β-carotene-derived apocarotenoids may contribute to improved seed longevity via a mechanism of regulating antioxidant enzyme gene expression and activities, although the exact mechanistic actions of apocarotenoids on them remain to be fully elucidated.

### Aquaporin TIP2;2 protein responds to β-cyclocitral signaling in improving seed longevity

Through comparative proteomics, we identified TIP2;2, a tonoplast intrinsic protein, which showed high abundance in *PSY* OE and low level in *psy* mutant seeds in comparison with WT. The *tip2;2* mutant seeds performed poorly in seed germination and accumulated high level of ROS after artificial aging. Both β-cyclocitral and β-ionone have been found to activate the promoter activity of *TIP2;2* in tobacco, indicating that apocarotenoids regulate its expression to contribute to seed longevity (Fig. 6).

Tonoplast intrinsic proteins represent a subfamily of the aquaporins superfamily, predominantly situated within the tonoplast (vacuolar membrane) of plant cells (Maurel et al. 2015). TIPs are essential for regulating water and solute transport across the vacuole, a process critical for cellular homeostasis and maintenance of turgor pressure for plant growth and development. Several studies have highlighted the role of TIPs including TIP2;2 in stress resilience (Sade et al. 2009; Wang et al. 2022) as well as in transporting ROS into the vacuole to contribute ROS detoxication within cells (Bienert et al. 2007; Wang et al. 2014).

Interestingly, Arabidopsis *TIP3;1* and *TIP3;2* have been shown to be involved in seed longevity. Seeds of the *tip3;1/tip3;2* double mutant accumulate high levels of hydrogen peroxide, whereas these *TIP* promoters are activated by ABSCISIC ACID INSENSITIVE 3 (ABI3) (Mao and Sun 2015), a key component in ABA-signal pathway which plays an important role in seed longevity (Liu et al. 2013). The *tip2;2* mutant seeds also accumulated high levels of ROS after aging, while the *TIP2;2* promoter responded to β-cyclocitral and β-ionone signaling (Fig. 6; Supplementary Fig. S10). It appears that *TIP2;2* contributes to seed longevity via activating by apocarotenoids and affecting ROS, although the precise molecular mechanisms underlying are unknown.

Notably, β-carotene is a metabolic precursor for abscisic acid (ABA) that greatly influences seed dormancy, desiccation tolerance, and germination (Shu et al. 2016). While these *PSY* genotypes were different in carotenoid levels and exhibited variations in seed longevity, they did not show any alterations in seed germination, implying less involvement of ABA in the *PSY* regulated seed longevity. Collectively, our data support that the β-carotene derived apocarotenoids play critical roles in improving seed longevity and in mediating *TIP2;2* expression among other unknown mechanisms.

## Materials and Methods

### Plant materials and growth conditions

The plant materials used included *Arabidopsis thaliana* (Columbia-0) wild type (WT) and T-DNA insertion lines (SALK_088744, SALK_076271, SALK_15463C) from Arabidopsis Biological Resource Center (ABRC). Transgenic Arabidopsis lines overexpressing tomato (*Solanum lycopersicum) PSY2* and *ccd1 ccd4* (*ccd1/4*) double mutant were as previously described (Cao et al. 2019; Dickinson et al. 2019). Plants were cultivated in soil in a walk-in growth chamber, which was set to provide 16 hours of light (150 µmol photons m^−2^ s^−1^) using T8 fluorescent lights, followed by 8 hours of darkness. The temperature was maintained at 23/20 °C day/night with 60% relative humidity. Tomato seeds of the Ailsa Craig (AC) cultivar and *psy1* mutant were germinated at 26/23 °C day/night with a 14/10 hrs light/dark cycle at Cornell’s Guterman Bioclimatic Laboratories. Seed samples for all the genotypes were collected after maturation at the same time.

### Seed germination assay

Both freshly collected seeds and control deteriorated seeds, as well as seeds after one-year natural aging in sealed bags at room temperature, underwent surface sterilization using 0.3% (w/v) sodium hypochlorite before being plated on half-strength MS medium. After a 3-day stratification period, seeds were germinated in growth chambers set at 22 ± 2 °C with a 16-hour light/8-hour dark cycle and a light intensity of 150 µmol photons m^−2^ s^−1^. Germination rates were assessed on the 7^th^ day. For the seed germination experiments, means and significance were derived from three biological replicates with at least 100 seeds per replicate. Seed germination was considered complete when the radicle extended beyond the testa and was evaluated until 7 days post-imbibition. Images of germinated seedlings were captured simultaneously for all the genotypes.

### Seed viability assay

Tetrazolium staining was conducted as described (Berridge et al. 2005). Surface-sterilized seeds were treated with 1% (w/v) 2,3,5-triphenyl tetrazolium chloride and incubated at 30 °C for 48 hours. Seeds were washed with water, and color changes were recorded following the reduction of non-color tetrazolium salt into a red formazan compound by viable seeds.

A spectrophotometric method was used to quantify formazan formation in the tetrazolium assay following the methods as described (Varshney et al. 2023). Seeds were incubated for 24 h in tetrazolium solution at 30 °C. Formazan production from 100 crushed seeds in 95% ethanol was measured for absorbance at 492 nm, using ethanol as a blank. Three biological replicates per genotype were analyzed.

### Controlled deterioration test (CDT)

The CDT is a common method for rapidly evaluating and predicting seed longevity and storability, which was carried out according to (Ogé et al. 2008). For the test, seeds were soaked in water for one hour and then incubated at 45 °C for 4 d for Arabidopsis seeds and at 50 °C for 5 d for tomato seeds at 100% relative humidity. After treatments, seeds were sterilized and plated out on half-strength MS medium under the same germination conditions described above for 7 days and germination rates were counted. To check the efficacy of β-carotene, lutein, β- cyclocitral, and β-ionone, WT Arabidopsis seeds were soaked with individual compounds at 5 μM, 10 μM, and 20 μM for 3 hours. CDT was performed, and germination was recorded. Experiments were conducted in triplicate, with each replicate containing approximately 100 seeds.

### The 3, 3’ - diaminobenzidine (DAB) staining

For DAB staining, seeds were incubated overnight at 25°C in the dark in the DAB staining solution (1 mg ml^-1^ DAB in 50 mM tris acetate buffer, pH 5.0). Thereafter, seeds were washed with 95% ethanol and photographed.

### Hydrogen peroxide (H_2_O_2_) and malondialdehyde (MDA) assays

For H_2_O_2_ assay, Arabidopsis seeds (50 mg) were ground with 2 ml of 0.1% trichloroacetic acid (TCA) and centrifuged at 13,000 g for 20 minutes at 4 °C. The supernatant (0.5 ml) was combined with 0.5 ml of 10 mM potassium phosphate buffer (pH 7.0) and 1 ml of 1 M potassium iodide in a total reaction mixture of 2 ml and incubated in the dark for 1 hour. The absorbance at 390 nm was recorded. H_2_O_2_ content was calculated using a standard curve created from known H_2_O_2_ concentrations.

MDA content was determined using the method as described (Tian et al. 2017). In brief, 50 mg of seeds were homogenized with 2 ml of 0.25% thiobarbituric acid in 10% TCA and incubated at 95 °C for 30 minutes. The reaction mixture was kept on ice for 10 minutes before centrifugation at 13,000 g for 30 minutes. The resulting supernatant was measured for absorbance at 532 nm and 600 nm. The MDA concentration in samples was calculated using an extinction coefficient of 155 mM^−1^ cm^−1^.

### Protein extraction

Seeds were crushed in liquid N2 and homogenized using an extraction buffer consisting of 100 mM HEPES (pH 7.5), 1 mM phenylmethylsulfonylfluoride, 1x protease inhibitor mixture (100x, Sigma), and 1 mM β-mercaptoethanol. The supernatant was collected after centrifuging at 10,000 g for 15 min at 4°C. Protein concentration was assessed employing the Bradford method (Bradford 1976).

### Catalase (CAT) and ascorbate peroxidase (APX) activity assays

Catalase activity was assessed using the method described (Boldrin et al. 2016). In the activity assay, 20 µl of total soluble seed protein extract was mixed with 3 ml of reaction mix containing 50 mM potassium phosphate buffer (pH 7.8) and 3 mM H_2_O_2_. After 2 min incubation, the decomposition of H_2_O_2_ by catalase was monitored at 240 nm for 60 s.

Ascorbate peroxidase activity was assessed following the method as described (Ramos et al. 2011). The complete reaction mixture included 50 mM potassium phosphate buffer (pH 7.8), 0.5 mM ascorbic acid (ASA), 0.5 mM H_2_O_2_, and 20 µl of seed extract in 1 ml of the reaction mixture. Ascorbate peroxidase activity was assessed by measuring the decrease in absorbance of ASA at 290 nm at 25°C, resulting from H_2_O_2_ oxidation for 3 minutes. The APX activity was calculated using 2.8 mM ¹ cm ¹ extinction coefficient.

### Gene expression analysis

Total RNA was extracted using Trizol reagent following the manufacturer’s protocol (ThermoFisher, Scientific, USA). The cDNA was synthesized using 4 µg of DNase I-treated RNA with MMLV Reverse Transcriptase (TaKaRa Bio, USA). RT-qPCR was performed on a CFX384 Touch Real-time PCR Detection System (Bio-Rad, USA) using SYBR Green Master Mix (Bio-Rad) and gene-specific primers (Supplementary Table S2) as detailed (Chayut et al. 2021). All RT-qPCRs were conducted with three biological replicates with three technical repeats.

### Carotenoid extraction and analysis

Carotenoids were extracted from seeds employing the method previously described (Gonzalez-Jorge et al. 2013; Sun et al. 2021). Briefly, approximately 100 mg of seeds were initially pulverized in liquid nitrogen, after which 50 mg of the ground materials were weighed and extracted with 450 µL of extraction buffer composed of methanol and chloroform in a 2:1 (v/v) ratio. The mixture was vortexed for 30 seconds, added with 450 µL of water and 150 µL of chloroform, and mixed thoroughly prior to centrifugation at 12,000 g for 10 minutes. The lower organic phase was then carefully transferred to a new 1.5 mL tube, dried using a SpeedVac, and redissolved in 100 µL of ethyl acetate.

Carotenoids were separated and quantified using ultra-performance convergence chromatography (UPC²). Chromatographic separation was performed on a Waters HSS C18 SB column (3.0 × 150 mm, 1.8 µm particle size) at 1.0 mL/min. Detection was carried out using a photodiode array detector, monitoring wavelengths from 250 to 700 nm. The column was equilibrated with 99% eluent A (supercritical CO) and 1% eluent B (methanol). Carotenoids were eluted using a nonlinear, concave gradient that ramped to 20% B over 7.5 minutes, held for 4.5 minutes, then returned to 1% B, followed by 3 minutes of re-equilibration. Individual carotenoids were identified by comparison to authentic standards (Sigma) and quantified using MassLynx software v4.1 (Waters) at compound-specific wavelengths chosen for optimal sensitivity and selectivity. A five-point β-carotene calibration curve (5–100 ng/mL) was used for quantification, and all concentrations are reported as β-carotene equivalents.

### Western blot analysis

For western blot analysis, 50 μg total proteins were separated using 15% SDS-PAGE, transferred onto a nitrocellulose membrane (Cytiva, Amersham), and stained with Ponceau S to evaluate protein loading. The membrane was then incubated with various protein-specific antibodies as previously outlined (Sun et al. 2023). Protein signals were detected utilizing the WesternBright ECL kit (Advansta, USA).

### Plant transformation

To overexpress *SlPSY2* in the *ccd1 ccd4* double mutant, the coding sequence (CDS) without a stop codon was amplified using gene-specific primers (Supplementary Table S3) and cloned into the pCAMBIA vector (Zhou et al. 2015). The construct was introduced into *Agrobacterium tumefaciens* GV3101 by electroporation and transformed into Arabidopsis *ccd1 ccd4* mutant using the floral dip method. Positive transgenic lines were selected through western blot analysis with an anti-PSY antibody at a dilution of 1:500. Three independent T3 homozygous transgenic lines were selected and utilized for the study.

### Agrobacterium-mediated infiltration in *N. benthamiana* leaves

Agrobacterium cells GV3101containing *SlPSY2* overexpression construct were cultured in LB medium supplemented with appropriate antibiotics until they achieved an absorbance of 0.6 at 600 nm. The bacterial cells were then pelleted, washed, and resuspended in an infiltration medium consisting of 10 mM 2-(N-morpholino)ethane sulfonic acid, 10 mM MgCl_2_, and 0.15 mM acetosyringone, adjusted to pH 5.6. This suspension was incubated at 25 °C for 4 to 6 hours to 0.6 OD at 600 nm before being infiltrated into the leaves of 4-weeks old *N. benthamiana* plants. The infiltrated plants were subsequently kept in the dark at 24 °C overnight, followed by a light cycle of 16 hours of light and 8 hours of darkness. After 48 hours, fluorescence in the leaf samples was examined using a laser scanning confocal microscope (Leica TCS-SP5) equipped with an argon laser and a 20× excitation source (Rao et al. 2024a).

### Comparative proteomic analysis

Proteomic analysis was performed as detailed (Supplementary Methods S1). Total proteins from seeds of WT, *PSY* OE and *psy* mutant lines with three biological replicates were extracted following the method described (Verma et al. 2013). For trypsin digestion and TMT 18-plex labeling, 25 µg of proteins for each sample were first reduced with Tris (2-carboxyethyl) phosphine (TCEP) and cysteines were blocked with methylmethanethiosulfonate (MMTS) and then digested. The resultant peptides were labeled before High pH Reverse Phase (hpRP) chromatography. The fractionated samples were subjected to NanoLC-MS/MS Analysis. Raw MS spectra were processed and searched against the Arabidopsis Araport11 database (https://www.uniprot.org/). The proteomics data were deposited to the ProteomeXchange Consortium with the dataset identifier (#PXD063818).

### Dual reporter assay

The pDual reporter assay was performed essentially as described (Sun et al. 2025). The pDual reporter construct with the *TIP2;2* promoter was transiently expressed in *N. benthamiana* leaves, which were treated with 5 and10 µM of β-cyclocitral or β-ionone and allowed to grow for 24 hours under a 16 h light and 8 h dark cycle. The leaves were detached, treated with a 0.1 mg ml^−1^ luciferin solution, and subsequently documented using the ChemiDoc MP system (BioRad).

To quantify tranactivation activity, total proteins from leaves were extracted and divided into equal volumes (100 µl) into an assay buffer containing either 1 mM luciferin or coelenterazine. FireflyLUC or nanoLUC was imaged using the ChemiDOC MP imaging system. The image intensity was quantified following conversion into grayscale utilizing IMAGEJ (Schneider et al. 2012). The FireflyLUC activity was used to normalize for the transactivation activity. Three replicates were carried out using three leaves of *N. benthamiana*.

### Statistical analysis

All data in this study are shown as means ± SD with three biological replicates. Statistical analysis was conducted using a one-way ANOVA. Duncan’s Multiple Range Test (*p* = 0.01) was performed using the SPSS program (SPSS, Chicago, IL, USA) for statistical significance.

## Supplementary data

The following materials are available in the online version of this article:

---

**Supplementary Figure S1.** Gene structure of *PSY* and the relative expression of *PSY* in *psy* mutants.

**Supplementary Figure S2.** Seedling vigor test of freshly harvested seeds of WT, *PSY* OE, and *psy* mutant.

**Supplementary Figure S3.** Seedling vigor test of WT, *PSY* OE, and *psy* mutants after natural and artificial aging.

**Supplementary Figure S4.** *PSY* expression affects seed longevity during natural and artificial aging in tomato.

**Supplementary Figure S5.** Relative expression levels of antioxidative genes in the seeds of WT, *PSY* OE, and *psy* mutant lines.

**Supplementary Figure S6.** Immunoblot analysis of PSY protein before and after controlled deterioration test (CDT) in Arabidopsis seed.

**Supplementary Figure S7.** Relative expression levels of *AtCCD4* and *AtCCD1* in seeds of WT and three *PSY* ccd1/4 lines.

**Supplementary Figure S8.** Seed germination and tetrazolium staining for viability test of WT, *PSY* OE, *ccd1/4,* and three *PSY ccd1/4* lines in control and after artificial aging.

**Supplementary Figure S9.** Seed germination of WT seeds treated with D-15 in control and after artificial aging and *β*-cyclocitral treatment of one-year naturally aging *psy* mutants.

**Supplementary Figure S10.** Gene structure of *TIP2;2* and relative expression.

**Supplementary Figure S11.** Transactivation assay of *TIP2;2* promoter by β-ionone.

**Supplementary Table S1.** Total proteins identified in proteomic analysis

**Supplementary Table S2.** List of differential abundance proteins in proteomic study.

**Supplementary Table S3.** List of primers used in this study.

**Supplementary Methods S1.** Detail method of proteomic analysis.

## Acknowledgments

We thank Dr. Alexandra Jazz Dickinson for providing D-15 and *ccd1ccd4* double mutant. We thank Cornell Gutterman Greenhouse and the Boyce Thompson Institute for providing the plant growth facility.

## Author Contributions

A.H. and L.L. designed and conceptualized this work. A.H. performed the majority of the experiments and analyzed the data. S.R. and M.J.C. assisted in growing and harvesting of Arabidopsis and tomato seeds and with some experiments. X.Z. and E.W. generated *PSY ccd1 ccd4* T1 seeds and/or helped in growing *Nicotiana* and aided some experiments. Y.Y. and T.F. performed proteomic analysis. T.F. aided carotenoid analysis. T.T., J.L., and MM contributed research agents, assisted with data interpretation, and revised the manuscript. A.H. and L.L. wrote the manuscript with contributions from all coauthors.

## Data availability

The data that support the findings of this study are available from the corresponding authors upon request.

