## Supplemental data for "*Phytoene synthase* modulates seed longevity via the action of β-carotene derived metabolites"

### Slide 1
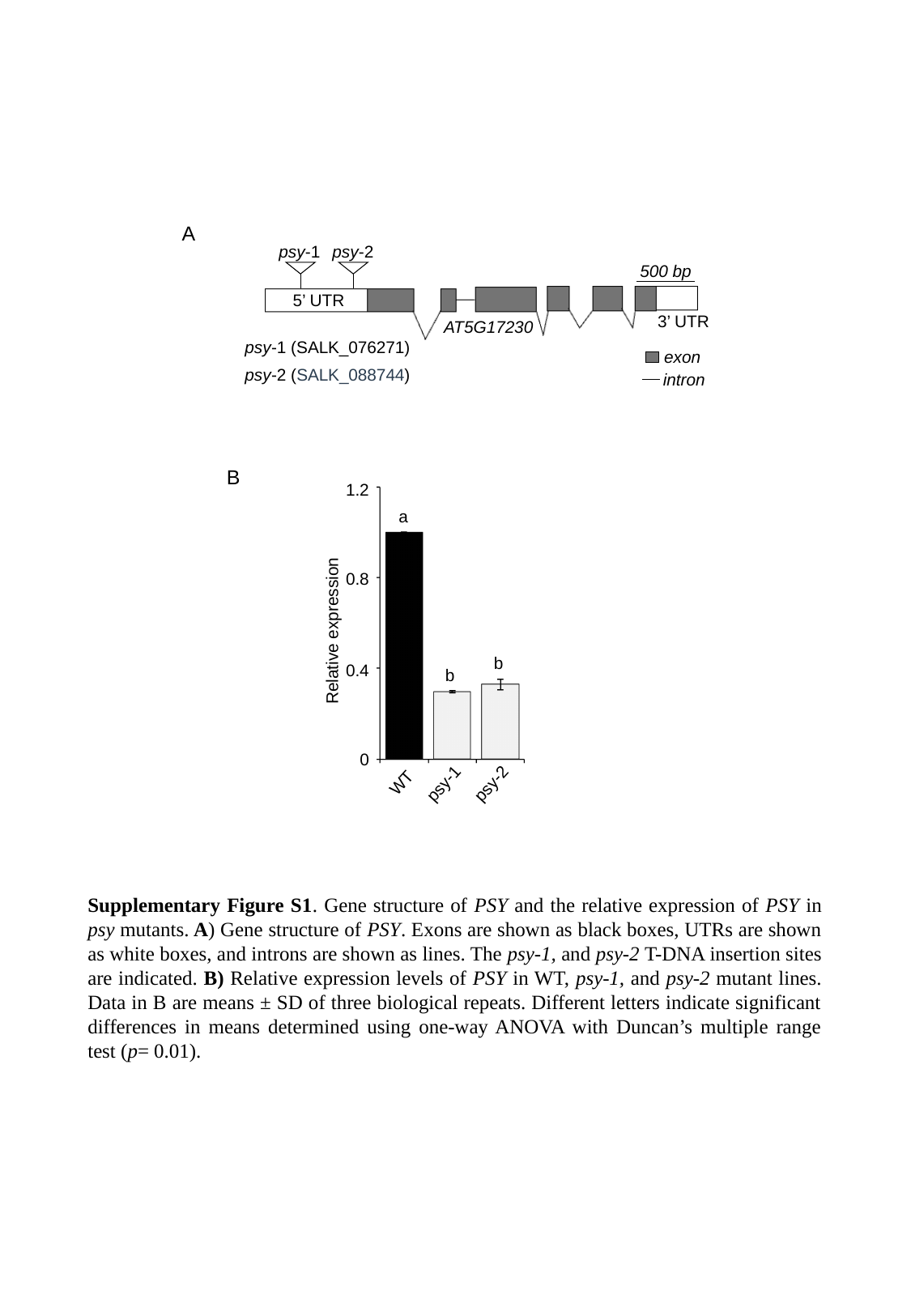

C T C G A G A G T C G C A T C T C G C C A G T C A A A A A C T A A A T T A A A A
41 A A C T A T A T A G A C C C A A A A A A T A A A A A T A A C G A C T T T T G T A
81 T T T A T A C T G G C A A G A T T C T C A T T A A A A C C C A G C A A A A A A A
121 A A A A A A A A A G A C A A A G A A G G A A G A A A A T T T G A G T G G G T G A
161 G A A T T T T T T A C G A T A G A G G A A G A G A C A G C A T C A T C T A C T T
201 G T G T T G T C T G T G T A C A T A T A T T A C A G T A A G C G T T G C A A T A
241 T A A C T T C T T G A G G A T C T T C T C A C A T T A A T G G G T C A A A C C T
281 T T T G C T C T T C C T T T T G A T T A A T T T A G T G T T T G A C A A T C T C
321 C T C C T C C T T C T C C T T C T T C T T C A A A G T T T T G T C G C A G T A T
361 C T A T T G T T C T T A C A G A G A G A A A G G T A G G C T T T T G T G T C C A
401 T C A C T C A T T T T C G C T C G T T T G G T T A A G C T T C A T C T G C C A T
441 G T G G T T C A C T G T T T T G A T A C T T T T G G G C T C T T A T A C C T A A
481 T G T T G A T G T A A C A C A A C G A T A C A T A A T C T A A T T T G T T T T C
521 G A G T G G A A A A T A G T T G A A C C C A G T T G A G T A T T T A G C C A A T
561 A C G A T T A G A G G A T T T A C T T G T T T T T T C T C T T T T C A T G T T C
601 T T C A T T T G C T T T T G A A G A T C T T C C T C T T T A T G T T T G T G T C
641 T C T T C G C T A T T T T A C T T G C T T G A A G A A A T T G A G T T T C T C A
681 T C T C G C T T A A A T T T G G C T T T A T A T A C A G A T T T A G A G A T C T
721 C G A T T C C T A A T C T A C T T G T T T T T G T A T T C A A T T T G C A G G A
761 A A G C T T T A G T C T T T T A C C A G T T T G A T C C A A T T C T G G G T T T
801 C A C T G A A A A A A A G T T G G G A G T T T G A T T C T T C T A A C T G T A G
841 A A G A A A C A G A G T C A A C A G A A G A A A A C T A A A A A A G T T G A G A
881 T T T T T C T C T C A C G C G C T C A A G A A C T T G A G T A T G T C T T C T T
921 C T G T A G C A G T G T T A T G G G T T G C T A C T T C T T C T C T A A A T C C
961 A G A C C C A A T G A A C A A T T G T G G G T T G G T A A G G G T T C T A G A A
1001 T C T T C T A G A C T G T T C T C T C C T T G T C A G A A T C A G A G A C T A A
1041 A C A A A G G T A A G A A G A A G C A G A T A C C A A C T T G G A G T T C T T C
1081 T T T T G T A A G G A A C C G A A G T A G A A G A A T T G G T G T T G T G T C T
1121 T C A A G C T T A G T A G C A A G T C C T T C T G G A G A G A T A G C T C T T T
1161 C A T C T G A A G A G A A G G T T T A C A A T G T T G T G T T G A A A C A A G C
1201 T G C T T T G G T G A A C A A A C A G C T A A G G T C T T C T T C T T A T G A C
1241 C T T G A T G T G A A G A A A C C A C A A G A T G T T G T T C T T C C T G G G A
1281 G T T T G A G T T T G T T G G G T G A A G C T T A T G A T C G A T G C G G T G A
1321 A G T T T G C G C T G A A T A T G C T A A G A C G T T T T A T C T T G G T A T G
1361 G A T C T T T T T A A C T C T T C T T T G G C T T T G G T G T A A T G T A G A G
1401 T G G T G T T C T G A G T T C T A A T G T T T G T G T T G C A G G A A C T T T G
1441 C T T A T G A C A C C C G A A A G G C G A A A G G C G A T T T G G G C A A T C T
1481 A C G G T A A G T T A C T G C A C A A A A C C A A T G G T T G A A G A G C T G T
1521 T T T A G A A T A T T T G C A T T G T C A A C A T T G A T T A T A A T T G T A G
1561 G A T A A A G T G A T G C T C A A A G T A G A T T T C T A C A A A C A A T C T A
1601 T T G T G G C T C T T G G T T A G T A T C T A G T T A A G T T G T C T T A A C A
1641 A G A G A A G T T G A A G C T G A A A T A G A T T C C T T G A A T C T T A C A C
1681 T C A T G T A G A A A G A T A G G T C T C T A G T T A G T T T G T G T G G T T A
1721 C C T T G T G T C T A A G A T A A G T C C A T G A A C A C C A A G C A T C C A A
1761 A T T T G C A C C C T T T G A A G A T A A A G A A T A A A G A C C C A A A A A G
1801 A C T A G C T T A G T T A C T A A T A T G G G T T T G T C C A A A A A A A A A A
1841 A A G T T A C T A A T A T G G G T T T T T A A T G G G C T T T T T T G G C C C A
1881 A T G G T C T T A A A C T C T G G A G A T T T T A C T C T A A T G A C C C C C T
1921 G T C T A G G A C A T A C T A G T A T C C A A A G C A T A T G G G G A A T T T T
1961 A A A G T T G T T T T C T T A T A T G T T T G A T T G G T T T G A A T T T G A T
2001 A G T T T T A C T C T G T C T T T T G T G C A G T T T G G T G T A G A A G A A C
2041 T G A T G A A C T T G T G G A T G G G C C A A A T G C T T C A C A T A T A A C T
2081 C C C A T G G C T T T A G A T A G A T G G G A A G C A A G G T T A G A A G A T C
2121 T T T T C C G T G G T C G T C C T T T C G A T A T G C T T G A T G C T G C T C T
2161 C G C T G A T A C A G T T G C T A G A T A C C C G G T C G A T A T T C A G G T C
2201 A G C C T T G C T C T G T C T T A T C T T A C T T C C T C A T T T C A T T A C A
2241 T G A G T T A A G A G T T G C A T T C A G T C C A T T G C A A T A A A C C T T A
2281 T A A A A C T T T G G T C T A A T G T T T A T T T A T G T T G C A G C C A T T T
2321 C G A G A C A T G A T C G A A G G A A T G A G A A T G G A C T T G A A G A A A T
2361 C G A G A T A C C A G A A C T T C G A T G A T C T A T A C C T T T A C T G C T A
2401 C T A C G T C G C T G G A A C C G T C G G A T T G A T G A G C G T T C C G G T T
2441 A T G G G A A T C G A T C C T A A G T C G A A A G C A A C A A C C G A A A G T G
2481 T T T A C A A C G C T G C C T T G G C C C T T G G T A T A G C C A A T C A G C T
2521 T A C T A A C A T A C T C A G A G A C G T A G G C G A A G A G T G A G T C T T T
2561 T T A T A T A T T T T A A A T A G A G C T T T G G T T T C T T G A T T T C A C A
2601 T A T C T C A C T T A A A C C G G T T T C T T G A T T T C A C A T G C T T C T T
2641 T T A A A C C G G T T T C T T G A T T C A C A T A T C T C T C T T A A A T T G G
2681 T T T T G T T C T C A C T A T C A G T G C G A G A A G A G G A A G G G T T T A T
2721 C T G C C T C A G G A T G A A T T G G C T C A G G C T G G T C T T T C A G A T G
2761 A A G A C A T A T T C G C C G G A A A A G T A A C T G A T A A A T G G A G A A A
2801 C T T C A T G A A A A T G C A G C T T A A A C G A G C A A G A A T G T T C T T C
2841 G A C G A A G C T G A G A A A G G C G T C A C C G A G C T C A G T G C C G C T A
2881 G C A G A T G G C C T G T A A G T G T C T C T A A A A C A C T C T T C A G C C G
2921 C A A G T T T A C C G T T T A C T T A T T T C T T T T G G A T C G A T T T T A A
2961 C C C C G A G G A C T T G G C T G C A A T G T T T C A G G T A T G G G C T T C A
3001 T T G C T A T T G T A C A G G A G A A T A C T G G A C G A G A T T G A A G C G A
3041 A T G A T T A C A A C A A T T T T A C T A A G A G A G C T T A T G T G G G G A A
3081 A G T C A A G A A A A T T G C A G C T T T G C C A T T G G C T T A T G C T A A A
3121 T C A G T A C T A A A G A C T T C A A G T T C A A G A C T A T C G A T A T G A G
3161 A G C G A G A G G A A A G T G G A A C A A A A A C A A C C T A A G A G C G C T T
3201 T T T G T G A T T A A G A A A A A A C T T A G G C T C G A A T T T A T T A T G T
3241 T A A C T A A T A T A T A C A T A T T A A T G G G G A A G C A A A T T C T T A T
3281 A A T G T T A C A T T A T C T T T C T G A A T G T A A A A A A G T A T T T T T T
3321 T T T T T A A T T T T A T T T T A G T T A T A G T T C T C T T G G C C T A A T G
3361 A G G G G A T A T A C A A T T T T T G G T T G A T G A A A T A A T C T T G T G A
3401 T A T A C A T A C A G A A A G A A G T T C T T C T T A T A T T C G A A T A A T A
3441 G T A C
Powered by BioJS
A
psy-2
psy-1
500 bp
5’ UTR
3’ UTR
AT5G17230
psy-1 (SALK_076271)
exon
psy-2 (SALK_088744)
intron
B
1.2
a
0.8
b
0.4
b
0
psy-2
WT
psy-1
Relative expression
Supplementary Figure S1. Gene structure of PSY and the relative expression of PSY in psy mutants. A) Gene structure of PSY. Exons are shown as black boxes, UTRs are shown as white boxes, and introns are shown as lines. The psy-1, and psy-2 T-DNA insertion sites are indicated. B) Relative expression levels of PSY in WT, psy-1, and psy-2 mutant lines. Data in B are means ± SD of three biological repeats. Different letters indicate significant differences in means determined using one-way ANOVA with Duncan’s multiple range test (p= 0.01).

### Slide 2
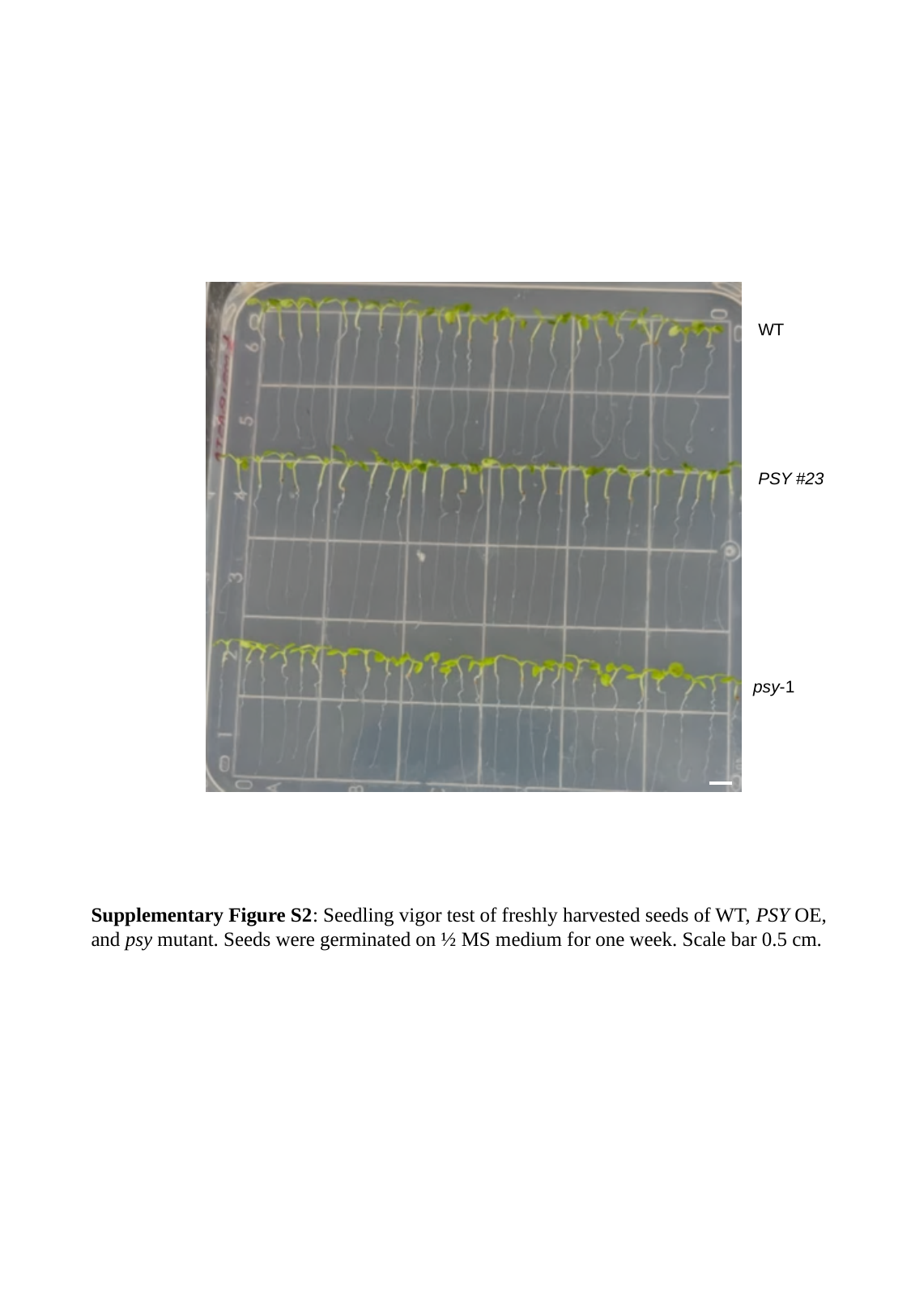

WT
PSY #23
psy-1
Supplementary Figure S2: Seedling vigor test of freshly harvested seeds of WT, PSY OE, and psy mutant. Seeds were germinated on ½ MS medium for one week. Scale bar 0.5 cm.

### Slide 3
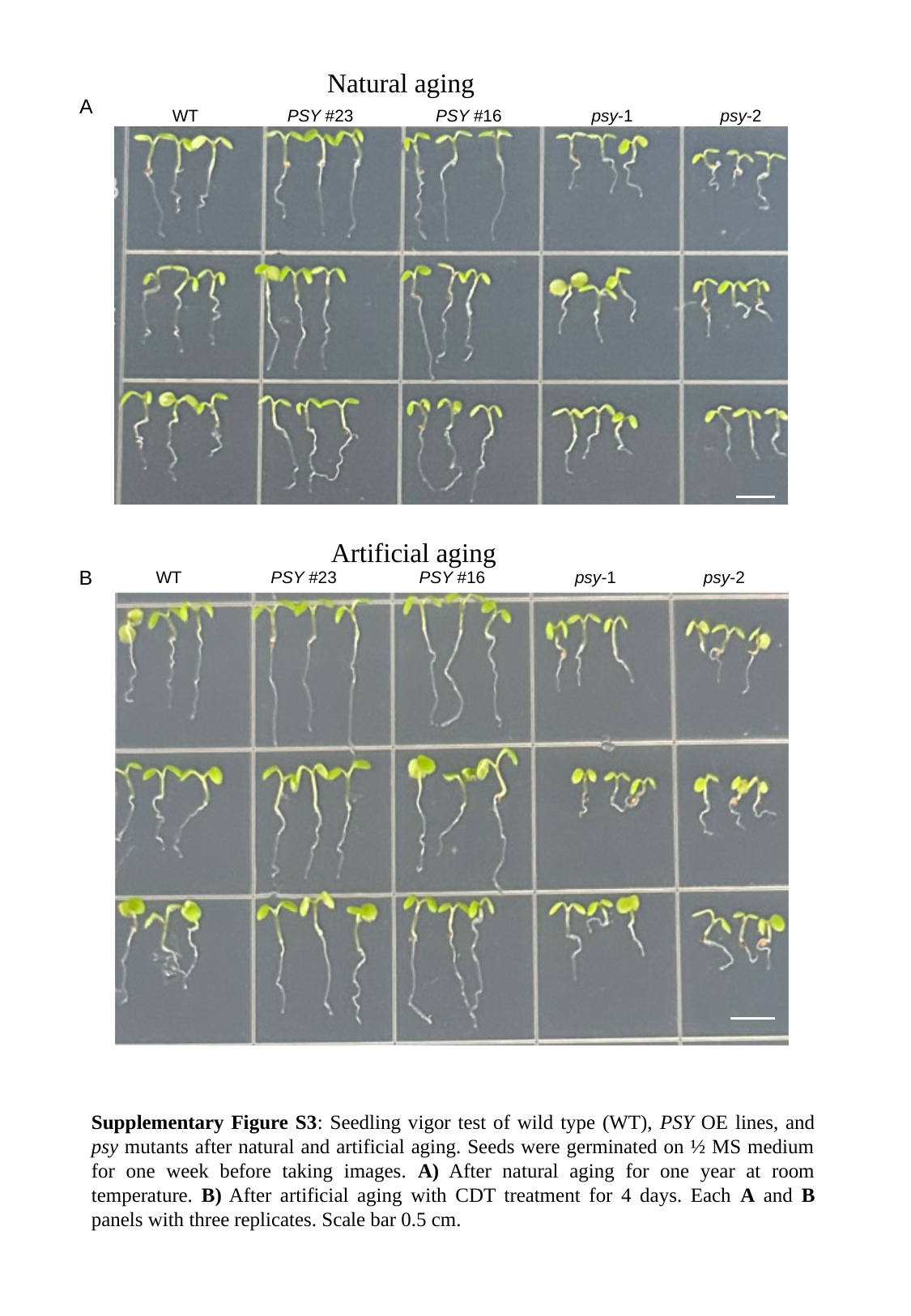

WT
PSY #23
PSY #16
psy-1
psy-2
Natural aging
A
Artificial aging
B
WT
PSY #23
PSY #16
psy-1
psy-2
Supplementary Figure S3: Seedling vigor test of wild type (WT), PSY OE lines, and psy mutants after natural and artificial aging. Seeds were germinated on ½ MS medium for one week before taking images. A) After natural aging for one year at room temperature. B) After artificial aging with CDT treatment for 4 days. Each A and B panels with three replicates. Scale bar 0.5 cm.

### Slide 4
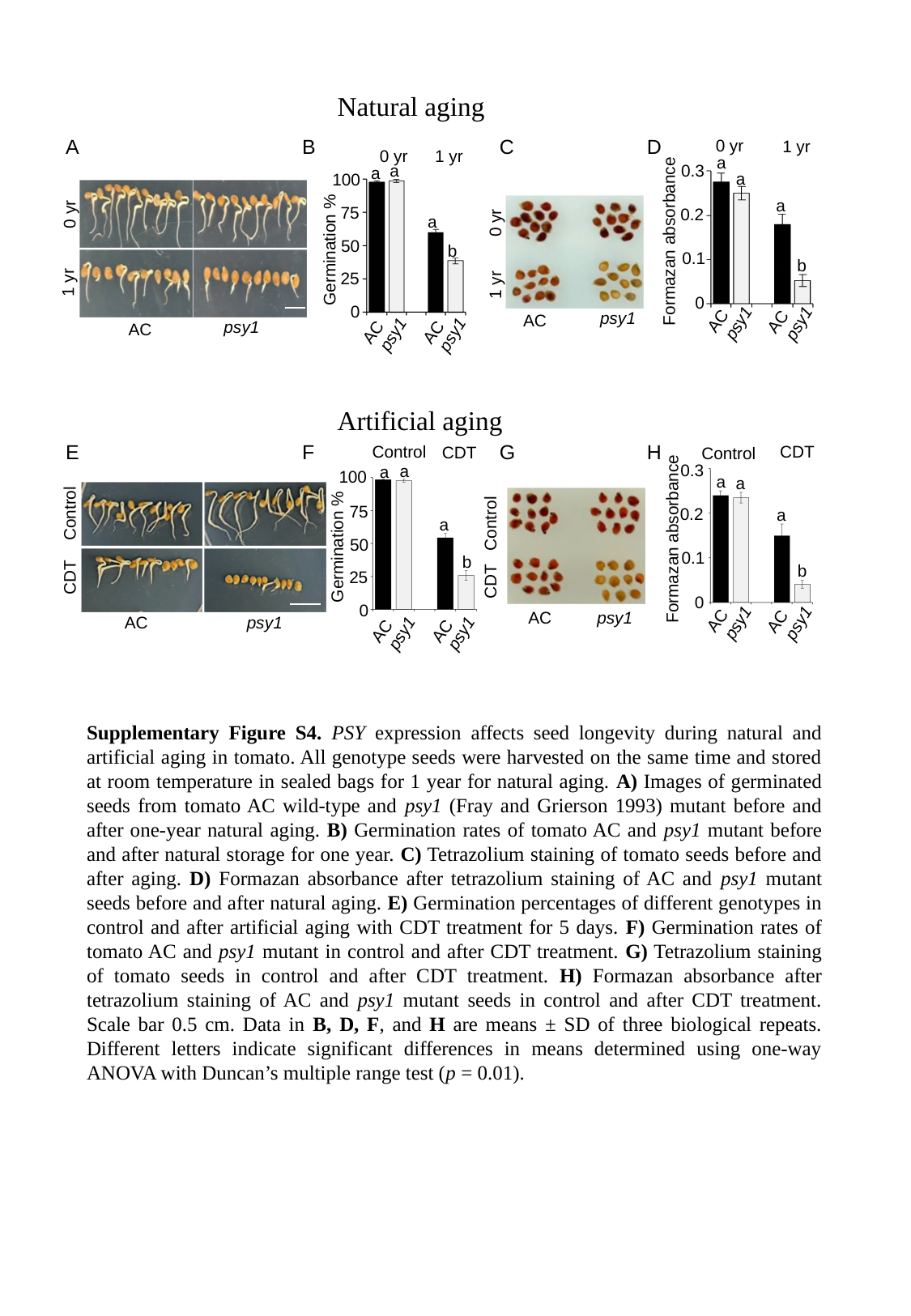

Natural aging
C
D
B
A
0 yr
1 yr
a
0.3
a
a
0.2
Formazan absorbance
0.1
b
0
AC
AC
psy1
psy1
1 yr
0 yr
a
a
100
75
a
50
Germination %
b
25
0
AC
AC
psy1
psy1
0 yr
1 yr
psy1
AC
0 yr
1 yr
psy1
AC
Artificial aging
G
H
F
E
CDT
Control
0.3
a
a
0.2
a
Formazan absorbance
0.1
b
0
AC
AC
psy1
psy1
Control
CDT
a
a
100
75
a
50
Germination %
b
25
0
AC
AC
psy1
psy1
Control
CDT
AC
psy1
Control
CDT
AC
psy1
Supplementary Figure S4. PSY expression affects seed longevity during natural and artificial aging in tomato. All genotype seeds were harvested on the same time and stored at room temperature in sealed bags for 1 year for natural aging. A) Images of germinated seeds from tomato AC wild-type and psy1 (Fray and Grierson 1993) mutant before and after one-year natural aging. B) Germination rates of tomato AC and psy1 mutant before and after natural storage for one year. C) Tetrazolium staining of tomato seeds before and after aging. D) Formazan absorbance after tetrazolium staining of AC and psy1 mutant seeds before and after natural aging. E) Germination percentages of different genotypes in control and after artificial aging with CDT treatment for 5 days. F) Germination rates of tomato AC and psy1 mutant in control and after CDT treatment. G) Tetrazolium staining of tomato seeds in control and after CDT treatment. H) Formazan absorbance after tetrazolium staining of AC and psy1 mutant seeds in control and after CDT treatment. Scale bar 0.5 cm. Data in B, D, F, and H are means ± SD of three biological repeats. Different letters indicate significant differences in means determined using one-way ANOVA with Duncan’s multiple range test (p = 0.01).

### Slide 5
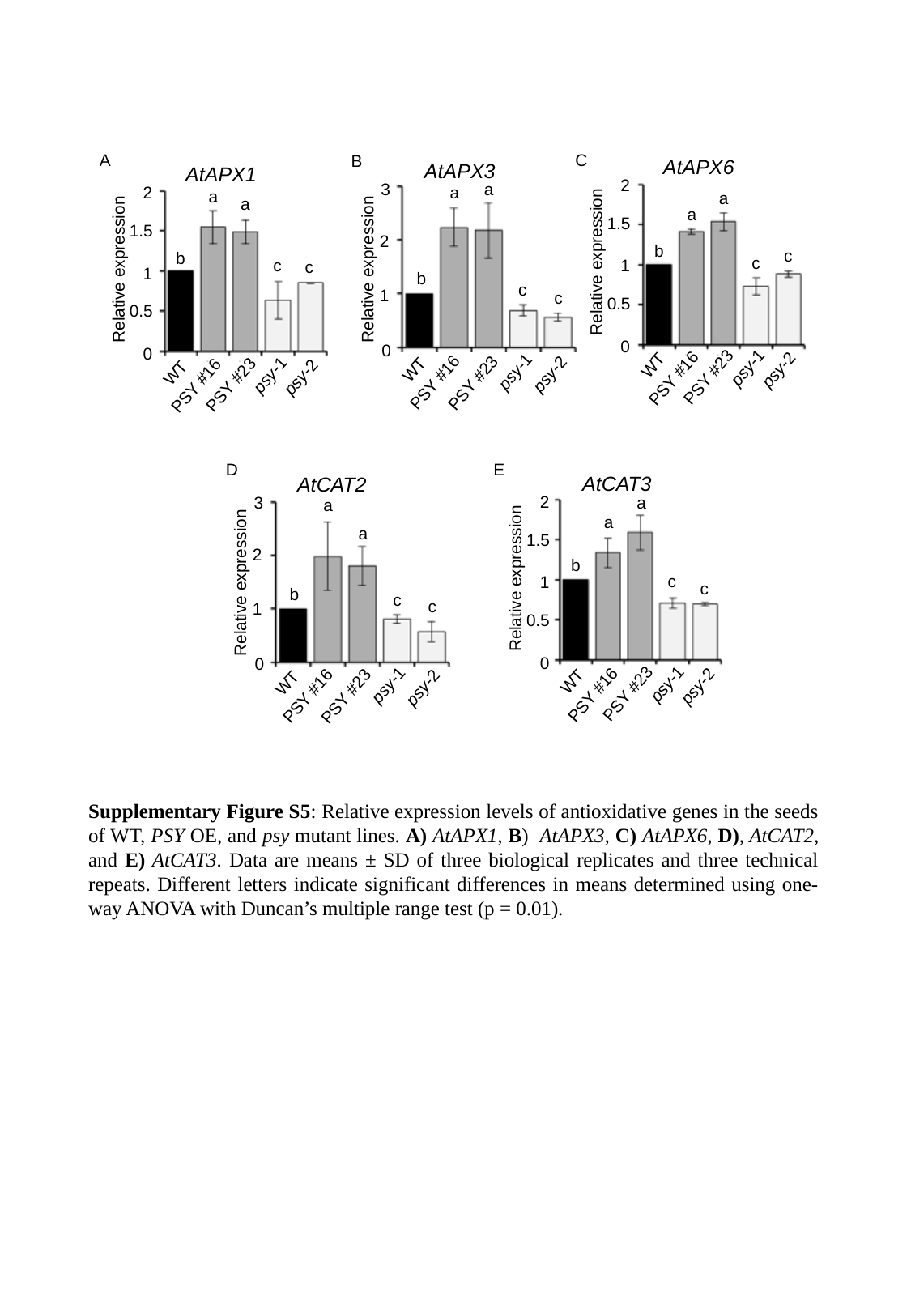

C
A
B
AtAPX6
2
a
a
1.5
b
c
c
1
Relative expression
0.5
0
WT
psy-1
psy-2
PSY #23
PSY #16
AtAPX3
a
a
2
b
Relative expression
c
1
c
0
WT
psy-1
psy-2
PSY #16
PSY #23
3
AtAPX1
2
a
a
1.5
b
c
c
1
Relative expression
0.5
0
WT
psy-1
psy-2
PSY #23
PSY #16
D
E
AtCAT3
2
a
a
1.5
b
c
1
c
Relative expression
0.5
0
WT
psy-1
psy-2
PSY #23
PSY #16
AtCAT2
a
a
2
b
Relative expression
c
c
1
0
WT
psy-1
psy-2
PSY #16
PSY #23
3
Supplementary Figure S5: Relative expression levels of antioxidative genes in the seeds of WT, PSY OE, and psy mutant lines. A) AtAPX1, B) AtAPX3, C) AtAPX6, D), AtCAT2, and E) AtCAT3. Data are means ± SD of three biological replicates and three technical repeats. Different letters indicate significant differences in means determined using one-way ANOVA with Duncan’s multiple range test (p = 0.01).

### Slide 6
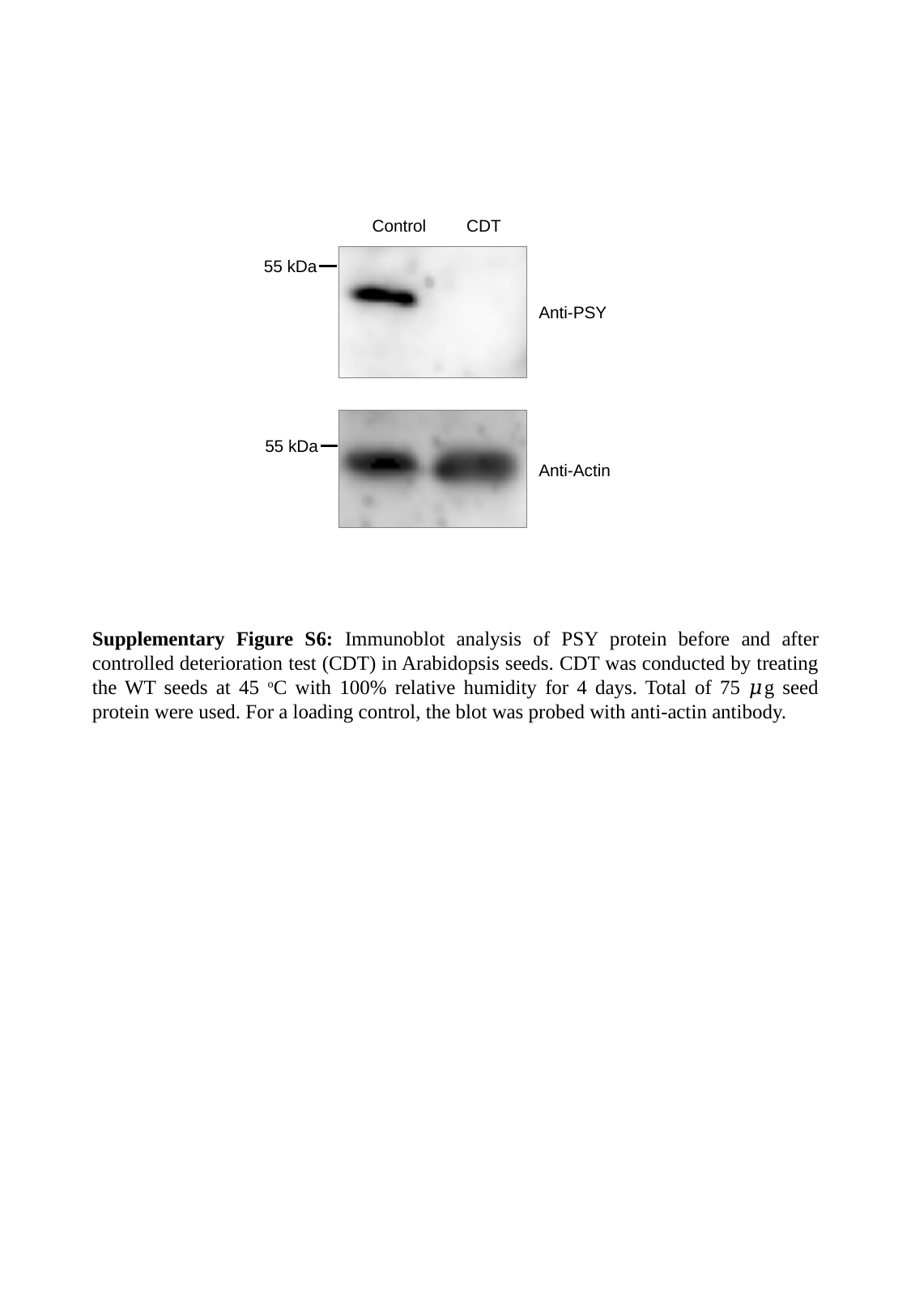

Control
CDT
55 kDa
Anti-PSY
55 kDa
Anti-Actin
Supplementary Figure S6: Immunoblot analysis of PSY protein before and after controlled deterioration test (CDT) in Arabidopsis seeds. CDT was conducted by treating the WT seeds at 45 ^o^C with 100% relative humidity for 4 days. Total of 75 𝜇g seed protein were used. For a loading control, the blot was probed with anti-actin antibody.

### Slide 7
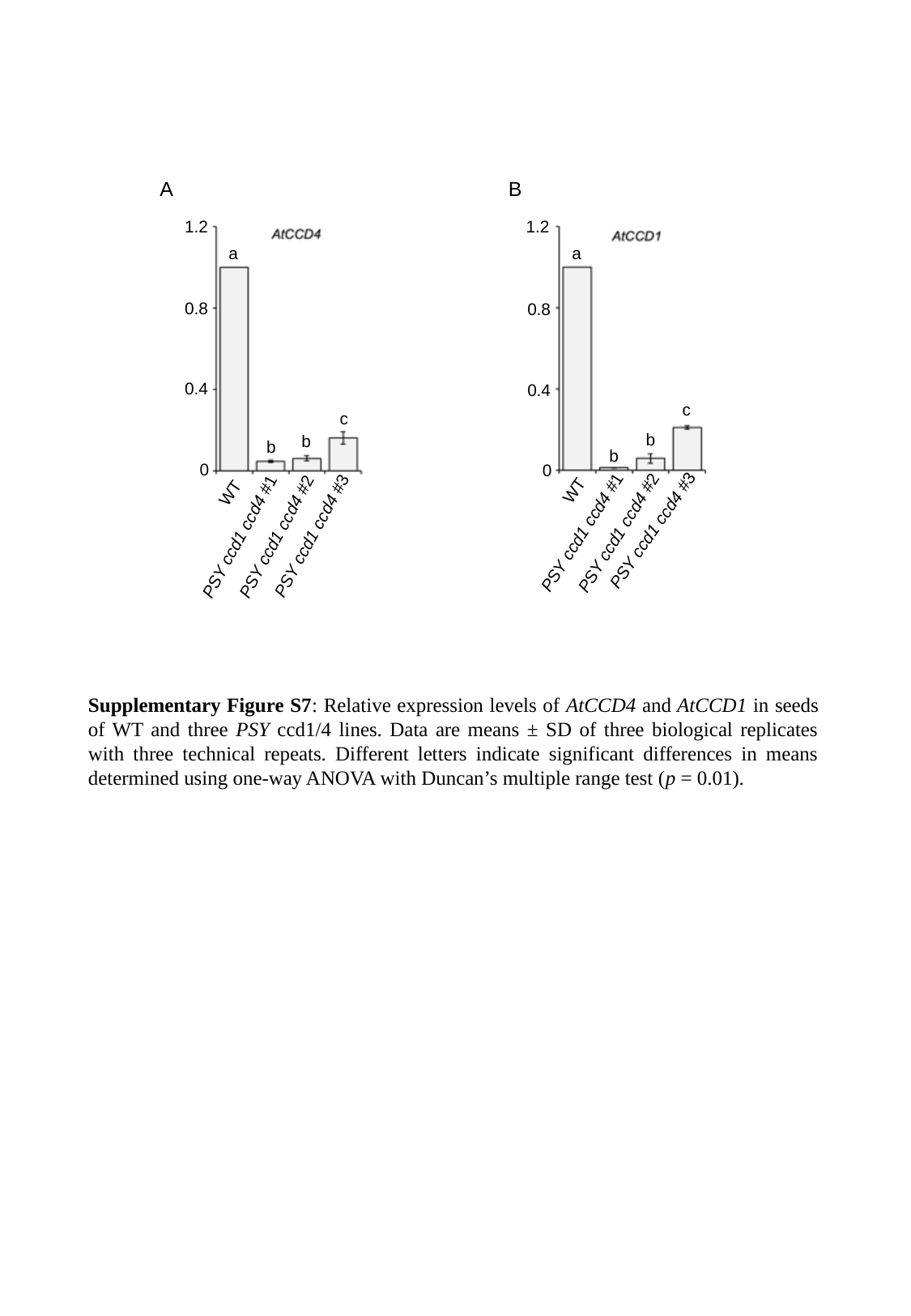

A
B
1.2
0.8
0.4
0
WT
PSY ccd1 ccd4 #3
PSY ccd1 ccd4 #1
PSY ccd1 ccd4 #2
1.2
a
0.8
0.4
c
b
b
0
WT
PSY ccd1 ccd4 #3
PSY ccd1 ccd4 #1
PSY ccd1 ccd4 #2
a
c
b
b
Supplementary Figure S7: Relative expression levels of AtCCD4 and AtCCD1 in seeds of WT and three PSY ccd1/4 lines. Data are means ± SD of three biological replicates with three technical repeats. Different letters indicate significant differences in means determined using one-way ANOVA with Duncan’s multiple range test (p = 0.01).

### Slide 8
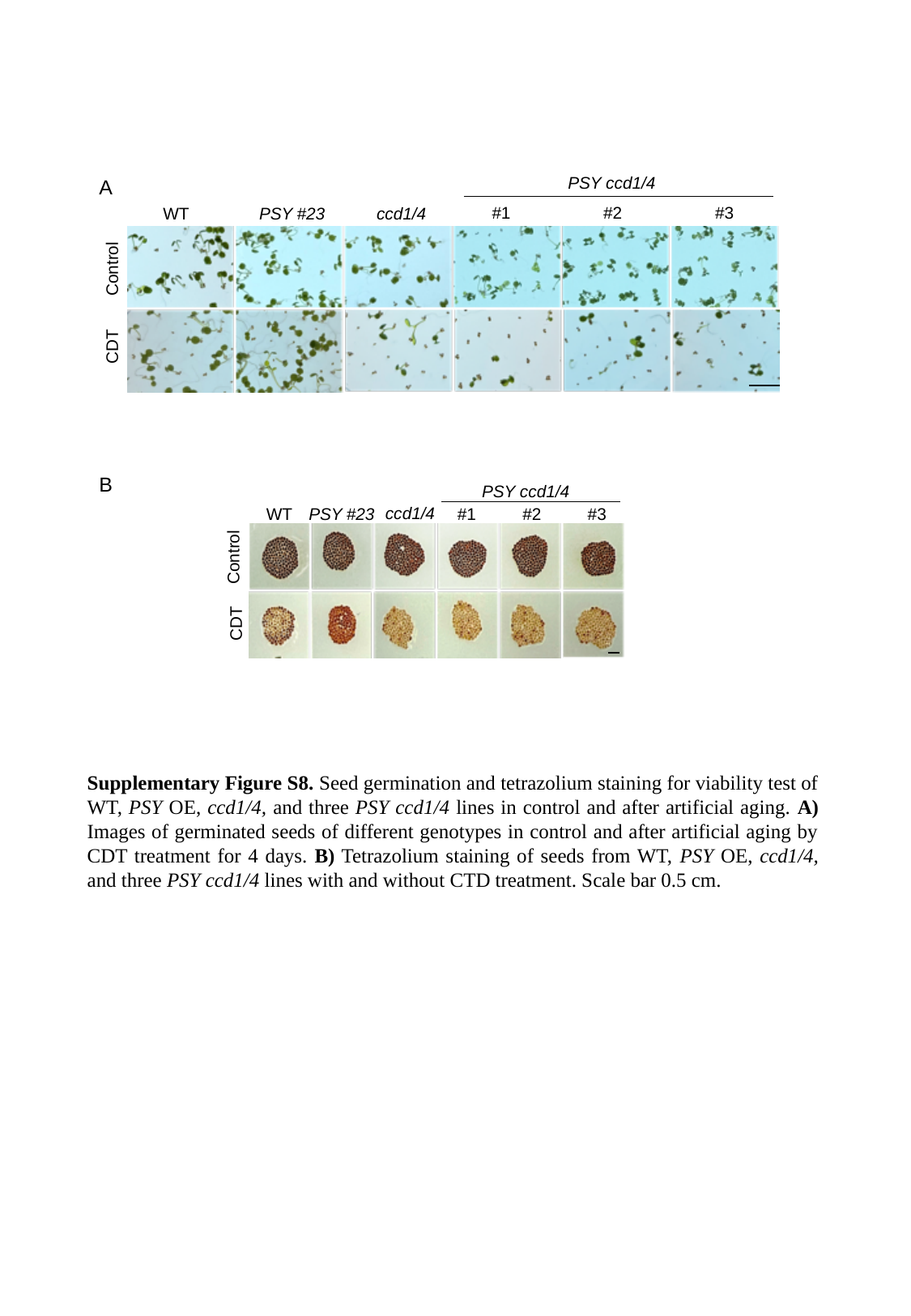

PSY ccd1/4
#1 #2 #3
WT
PSY #23
ccd1/4
Control
CDT
A
B
PSY ccd1/4
ccd1/4
WT
PSY #23
#1 #2 #3
Control
CDT
Supplementary Figure S8. Seed germination and tetrazolium staining for viability test of WT, PSY OE, ccd1/4, and three PSY ccd1/4 lines in control and after artificial aging. A) Images of germinated seeds of different genotypes in control and after artificial aging by CDT treatment for 4 days. B) Tetrazolium staining of seeds from WT, PSY OE, ccd1/4, and three PSY ccd1/4 lines with and without CTD treatment. Scale bar 0.5 cm.

### Slide 9
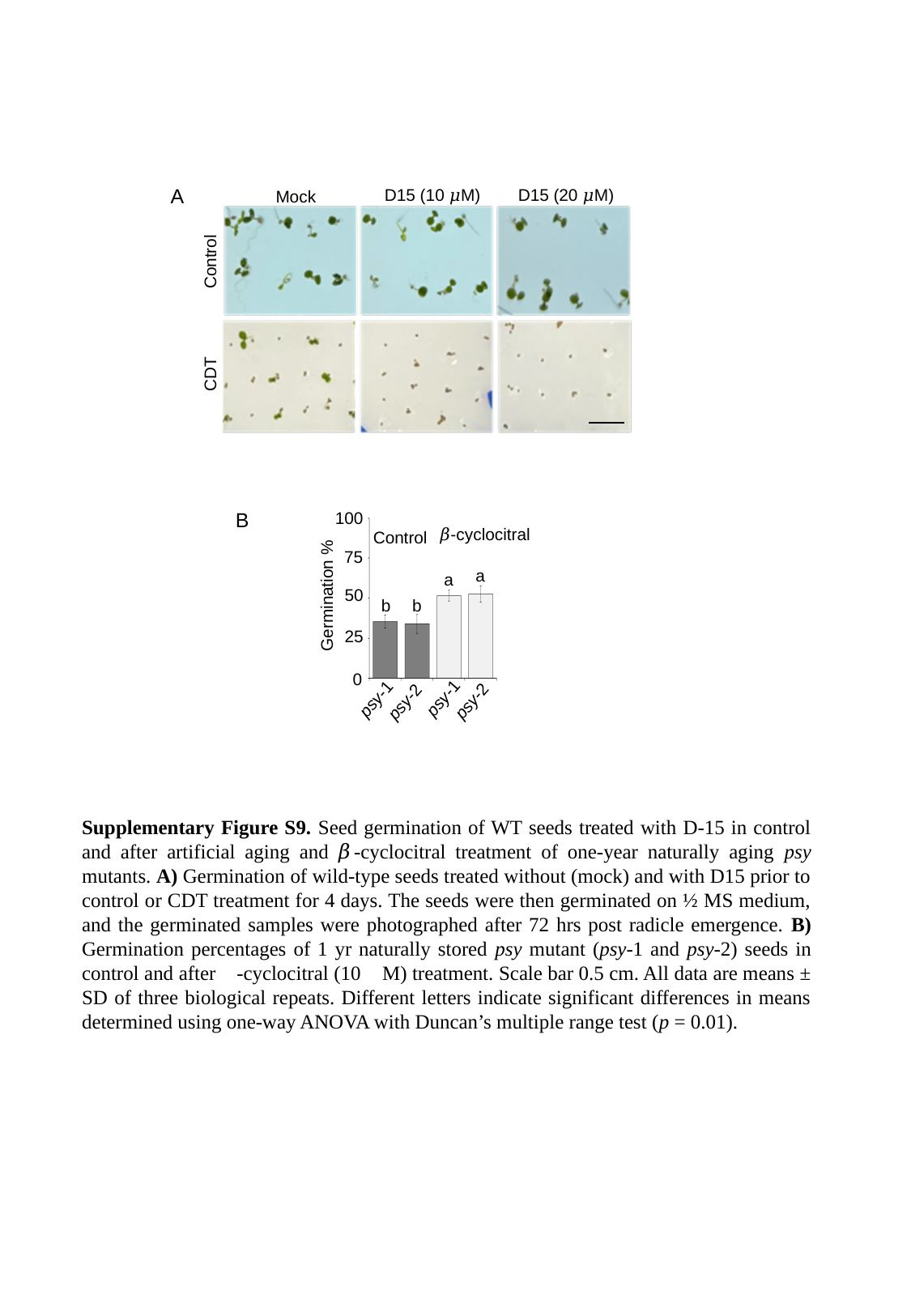

A
D15 (10 𝜇M)
D15 (20 𝜇M)
Mock
Control
CDT
B
100
75
a
a
50
Germination %
b
b
25
0
psy-1
psy-2
𝛽-cyclocitral
Control
psy-1
psy-2
Supplementary Figure S9. Seed germination of WT seeds treated with D-15 in control and after artificial aging and 𝛽-cyclocitral treatment of one-year naturally aging psy mutants. A) Germination of wild-type seeds treated without (mock) and with D15 prior to control or CDT treatment for 4 days. The seeds were then germinated on ½ MS medium, and the germinated samples were photographed after 72 hrs post radicle emergence. B) Germination percentages of 1 yr naturally stored psy mutant (psy-1 and psy-2) seeds in control and after 𝛽-cyclocitral (10 𝜇M) treatment. Scale bar 0.5 cm. All data are means ± SD of three biological repeats. Different letters indicate significant differences in means determined using one-way ANOVA with Duncan’s multiple range test (p = 0.01).

### Slide 10
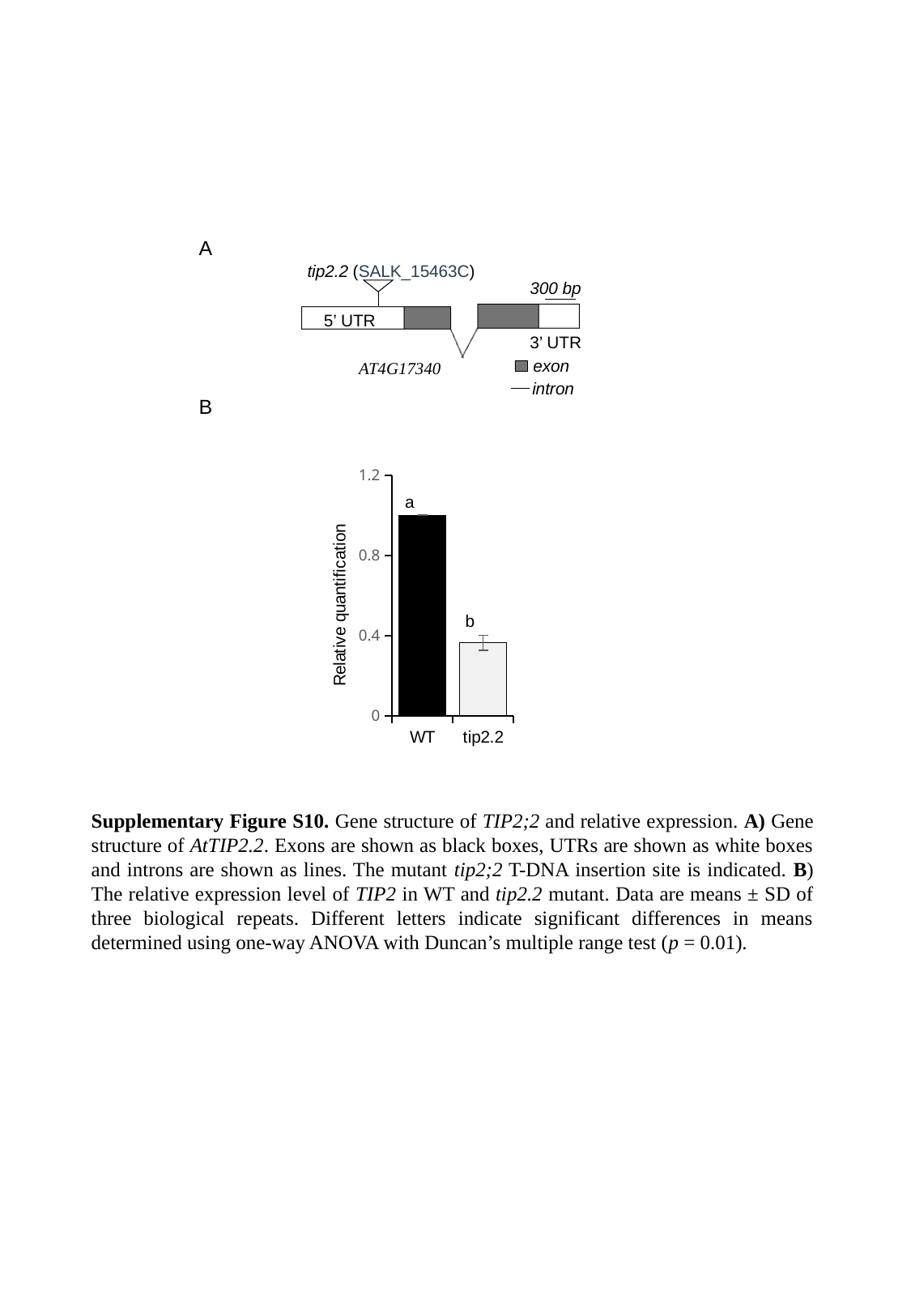

A
tip2.2 (SALK_15463C)
300 bp
5’ UTR
3’ UTR
exon
AT4G17340
intron
B
#### Chart
| Category | |
|---|---|
| WT | 1.0 |
| tip2.2 | 0.3629427198579873 |a
b
Supplementary Figure S10. Gene structure of TIP2;2 and relative expression. A) Gene structure of AtTIP2.2. Exons are shown as black boxes, UTRs are shown as white boxes and introns are shown as lines. The mutant tip2;2 T-DNA insertion site is indicated. B) The relative expression level of TIP2 in WT and tip2.2 mutant. Data are means ± SD of three biological repeats. Different letters indicate significant differences in means determined using one-way ANOVA with Duncan’s multiple range test (p = 0.01).

### Slide 11
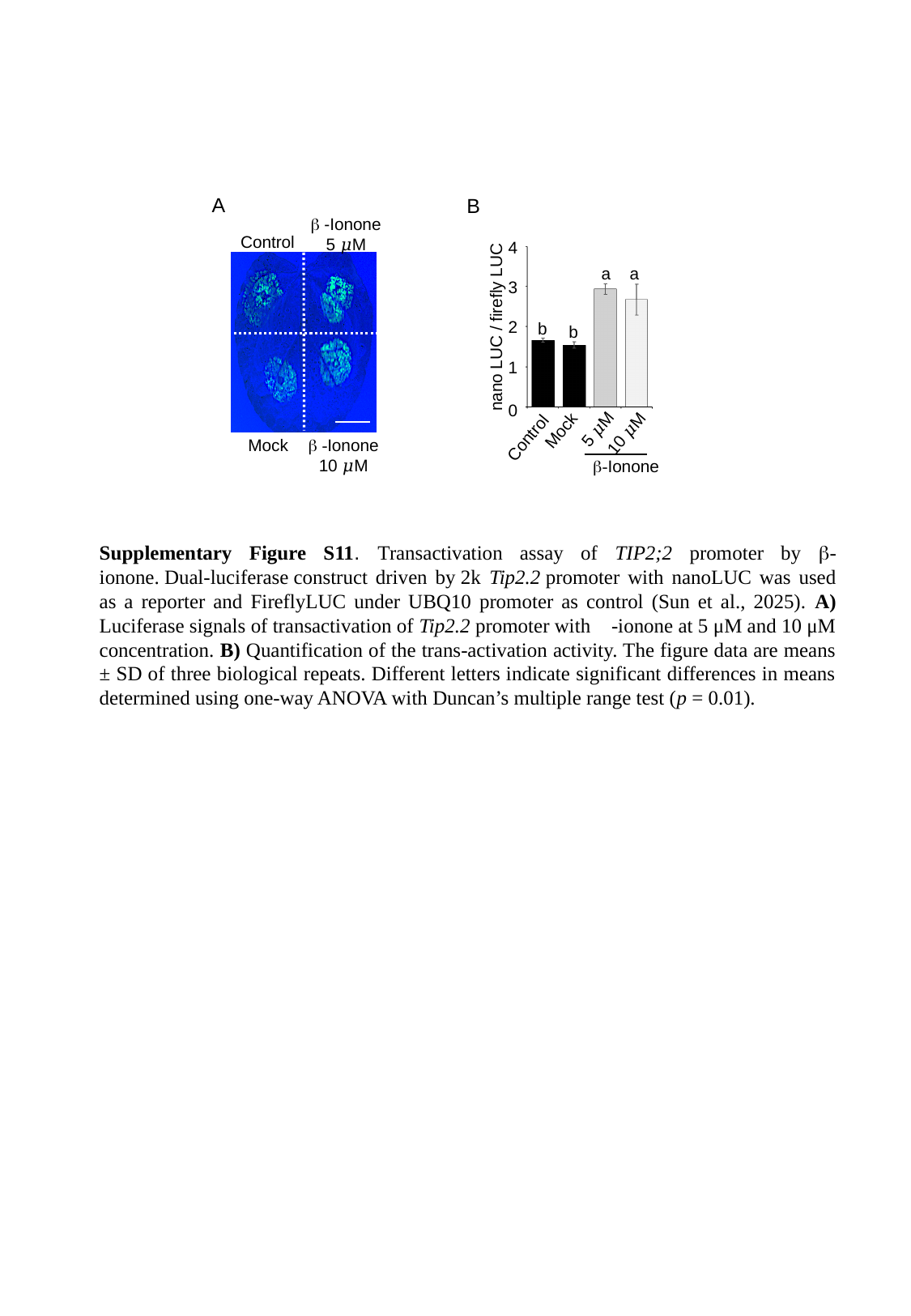

A
B
β -Ionone
5 𝜇M
Control
Mock
β -Ionone
10 𝜇M
4
a
a
3
2
nano LUC / firefly LUC
b
b
1
0
5 𝜇M
Mock
10 𝜇M
Control
β-Ionone
Supplementary Figure S11. Transactivation assay of TIP2;2 promoter by β-ionone. Dual-luciferase construct driven by 2k Tip2.2 promoter with nanoLUC was used as a reporter and FireflyLUC under UBQ10 promoter as control (Sun et al., 2025). A) Luciferase signals of transactivation of Tip2.2 promoter with 𝛽-ionone at 5 μM and 10 μM concentration. B) Quantification of the trans-activation activity. The figure data are means ± SD of three biological repeats. Different letters indicate significant differences in means determined using one-way ANOVA with Duncan’s multiple range test (p = 0.01).
